# The meadow jumping mouse genome and transcriptome suggest mechanisms of hibernation

**DOI:** 10.1101/2020.11.02.365791

**Authors:** Qian Cong, Ethan A. Brem, Jing Zhang, Jessica Alföldi, Jeremy Johnson, Elinor K. Karlsson, Kerstin Lindblad-Toh, Jason L. Malaney, William J. Israelsen

## Abstract

Hibernating mammals exhibit medically relevant phenotypes, but the genetic basis of hibernation remains poorly understood. Using the meadow jumping mouse (*Zapus hudsonius*), we investigated the genetic underpinnings of hibernation by uniting experimental and comparative genomic approaches. We assembled a *Z. hudsonius* genome and identified widespread expression changes during hibernation in genes important for circadian rhythm, membrane fluidity, and cell cycle arrest. Tissue-specific gene expression changes during torpor encompassed Wnt signaling in the brain and structural and transport functions in the kidney brush border. Using genomes from the closely related *Zapus oregonus* (previously classified as *Z. princeps*) and leveraging a panel of hibernating and non-hibernating rodents, we found selective pressure on genes involved in feeding behavior, metabolism, and cell biological processes potentially important for function at low body temperature. Leptin stands out with elevated conservation in hibernating rodents, implying a role for this metabolic hormone in triggering fattening and hibernation. These findings illustrate that mammalian hibernation requires adaptation at all levels of organismal form and function and lay the groundwork for future study of hibernation phenotypes.

## INTRODUCTION

Hibernating mammals exhibit extreme behavioral and metabolic phenotypes that generally include seasonal obesity and a winter-long fast, during which the hibernator employs repeated bouts of torpor^1^. During torpor, a near-freezing body temperature and greatly reduced metabolic rate promote energy conservation and survival during seasonal scarcity^2^. Mammals evolved from a heterothermic ancestor (i.e., one with variable body temperature) and torpor is an ancestral trait that has been retained in mammals ranging from marsupials to primates^3,4^. Torpor and other conserved hibernation phenotypes have untapped potential for application to human health^4^. Slowing metabolism, and thus reducing metabolic demand at the cellular level, could prolong viability of organs during storage for transplant, improve outcomes of medical emergencies involving insufficient blood flow or oxygen supply, and potentially enable prolonged spaceflight^5–7^. While significant progress has been made to understand the physiology of hibernation using multiple organisms including bears, bats, rodents, and lemurs, little is known about its genomic basis or molecular mechanisms^4^. Pursuit of hibernation mechanisms would be enabled by use of organisms that allow convenient genetic manipulation in the laboratory setting.

The meadow jumping mouse (*Zapus hudsonius)* is a small, hibernating North American rodent with a short generation time that provides an opportunity for genetic investigation of hibernation phenotypes^8,9^. Jumping mice accumulate significant fat stores in the pre-winter months and then hibernate without feeding for up to 9 months of the year^9–11^. During hibernation, their body temperature falls to near freezing and metabolic rate is reduced by as much as 98% while the animal is in a state of torpor^12,13^. Bouts of torpor are periodically interrupted by short interbout arousals, during which the animal rewarms for unknown reasons^1^. In meadow jumping mice, interbout arousals last for around 12 hours^8,13^ and torpor bouts vary in length throughout the hibernation season, but can last up to three weeks^14,15^.

Meadow (*Z. hudsonius*) and Great Basin jumping mice (*Z. oregonus*; previously western jumping mice *Z. princeps*) diverged from each other 2.5-3.7 million years ago^16,17^. While both species hibernate, they have adapted to use different strategies for controlling the timing of reproduction and hibernation. Meadow jumping mice depend heavily on photoperiod, such that long days are permissive for reproduction and prevent pre-hibernation fattening^8,18^; meadow jumping mice therefore produce two or three litters of young throughout the summer^19^ and then fatten in response to short days with cold temperature as a secondary cue^8,18^. In contrast, Great Basin jumping mice, which face shorter summers in a montane habitat, produce only one litter in the spring and then fatten based on increased food availability in late summer^20^. Great Basin jumping mice fatten whenever sufficient food is available, even when held in conditions of long photoperiod and warm temperature that would inhibit fattening in meadow jumping mice^20^.

To establish meadow jumping mice (*Z. hudsonius*) as a model for genetic studies of hibernation, we accustomed wild animals to laboratory environments^8^ and obtained genomes of multiple individuals. We induced fattening and hibernation in *Z. hudsonius* using controlled environmental cues, and analyzed the transcriptomes from a panel of seven organs and tissues during pre-hibernation fattening and torpor relative to lean ‘summer’ animals. Our data suggest that organ-specific gene expression changes occur to accommodate the torpor condition, including: i) down-regulated Wnt signaling in the brain possibly associated with lower synaptic activity; and ii) expression changes in the kidney to putatively adjust filtration function and maintain structural integrity in the cold. A small set of genes exhibited universal expression changes during torpor, suggesting more general adaptations including circadian rhythm (*Per3*), membrane fluidity (*Fads3*), and cell cycle arrest (*Cdkn1a/p21*) during hibernation.

We further generated additional genomes for Great Basin jumping mice (*Z. oregonus*). Comparative analysis of the two *Zapus* species revealed positively selected rapid divergence in the satiety hormone leptin, as well as differences in genes related to feeding behavior and sugar and lipid metabolism. When combined, these differences likely arise from the distinctive seasonality of food availability in their native environments and contribute to differential behavioral responses to environmental cues controlling reproduction and hibernation. Finally, we expanded our genome-genome comparison to eleven mammalian genomes and identified proteins showing significantly greater conservation in hibernating rodents compared to non-hibernating mammals. Notably, the membrane fluidity modulator Fads3 and the hormone leptin that stand out in comparisons of jumping mice also show elevated conservation across hibernators, suggesting their important role in hibernation. The work reported here provides insight into the genetic basis of hibernation phenotypes and lays the groundwork for the use of the meadow jumping mouse as a laboratory model of hibernation.

## RESULTS AND DISCUSSION

### *Inducing hibernation in* Z. hudsonius *and chromosome-level assembly of its genome*

In lab conditions, we induced fattening and hibernation of *Z. hudsonius* by simulating winter with short photoperiod (8 hours light/16 hours dark) and cold temperature^8^. The meadow jumping mice fatten in response to these cues by increasing food consumption and they spontaneously enter hibernation once they have accumulated sufficient fat stores. In the active state, the meadow jumping mouse (Fig. 1a) maintains a stable, euthermic core body temperature, with minor temperature fluctuations that correspond to daily rhythms of sleep and activity (Fig. 1b). When hibernating, the meadow jumping mouse employs torpor, a state of inactivity during which core body temperature falls to match the cold environment (Fig. 1c)^21–23^.

**Figure 1.**
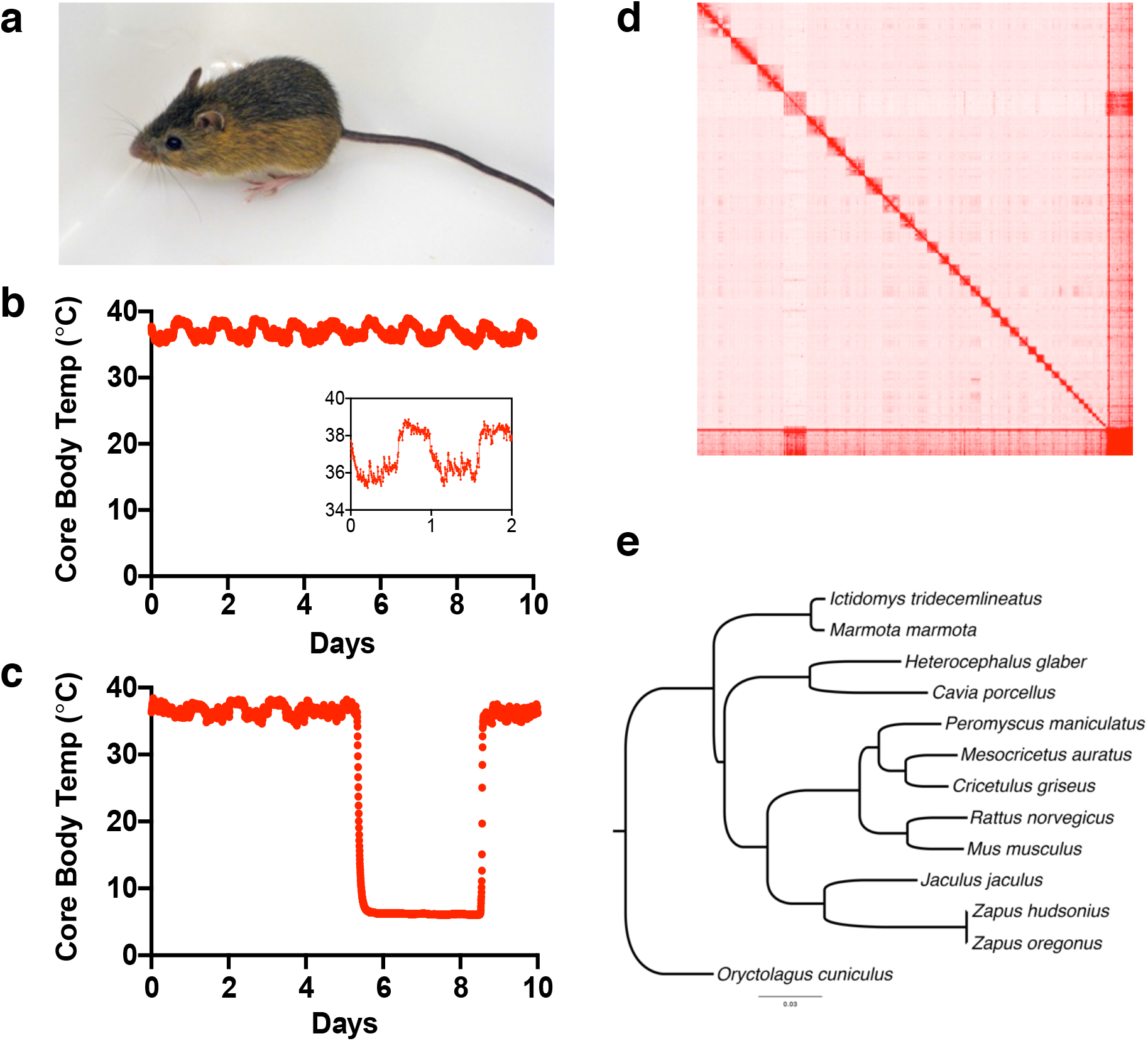
Meadow jumping mouse (*Z. hudsonius*). (**a**) Image of an adult meadow jumping mouse. (**b**) Core body temperature of a euthermic meadow jumping mouse is ~38°C and shows daily reduction of around 2°C during sleep (shown in inset). (**c**) Core body temperature falls to near 6°C ambient temperature during the first, short torpor of hibernation (shown here) and all subsequent torpor bouts. (**d**) The Hi-C contact matrix showing the meadow jumping mouse genome assembly. Each block of high contact represents one chromosome-scale superscaffold. (**e**) Rodent phylogeny generated from protein sequences of single copy orthologs, with rabbit (*Oryctolagus cuniculus*) as outgroup.

Using shotgun sequencing, Illumina mate-pair libraries and Hi-C data, we obtained a 2.1 Gb assembly with 35 chromosome-sized superscaffolds (N50 = 50.8 Mb, Fig.1d). LASTZ alignment of the *Zapus* and *Mus musculus* genome assemblies shows long regions of synteny as well as chromosomal rearrangements, which is expected due to the different numbers of chromosomes between the two species. Seven of the 35 superscaffolds in *Z. hudsonius* predominantly (>75%) map to a single *Mus* chromosome (Table S1); 24 *Z. hudsonius* superscaffolds break into 2-3 chromosomes in *Mus*. While the fourth-largest *Zapus* superscaffold contains most regions aligning to the *Mus* X chromosome (Fig S1), three additional shorter superscaffolds also contain X-linked regions of *Mus* (Table S1).

Assisted by the transcriptomes of *Z. hudsonius* and homologs from other rodents, we annotated 21,522 protein-coding genes in the *Zapus* genome (Table S2). We found the orthologs for each gene from other rodents and used the single-copy orthologs to build the phylogenetic tree to place *Zapus* among other rodents (Fig. 1e). *Zapus* clusters with the lesser Egyptian jerboa (*Jaculus jaculus*) in the phylogenetic tree, which is consistent with the fact that they both are members of the family Dipodidae (Dipodoidea). Dipodid rodents share an ancestral relationship with the muroid rodents (Muroidea; clade containing mice, rats, gerbils and hamsters), thus they are more closely related to non-hibernating muroids than other hibernators such as the thirteen-lined ground squirrel and marmot from family Sciuridae.

### Changes in gene expression profiles during hibernation

To better understand the gene expression changes associated with pre-hibernation fat deposition and torpor, we performed mRNA sequencing and analyses on relevant tissues obtained during those conditions. We housed meadow jumping mice in simulated seasonal conditions, and obtained tissue samples from animals in a ‘summer’ active condition, animals that had fattened in the cold but not yet entered torpor (the ‘cold’ condition), and hibernating animals that had been torpid at 6°C for approximately 72 hours in their second or third torpor bout (the ‘torpor’ condition) (Fig. 2a). We sequenced tissues that we expected to be important for physiology during fattening and torpor, including brown adipose tissue (BAT), brain, heart, kidney, liver, skeletal muscle (gastrocnemius), and white adipose tissue (WAT).

**Figure 2.**
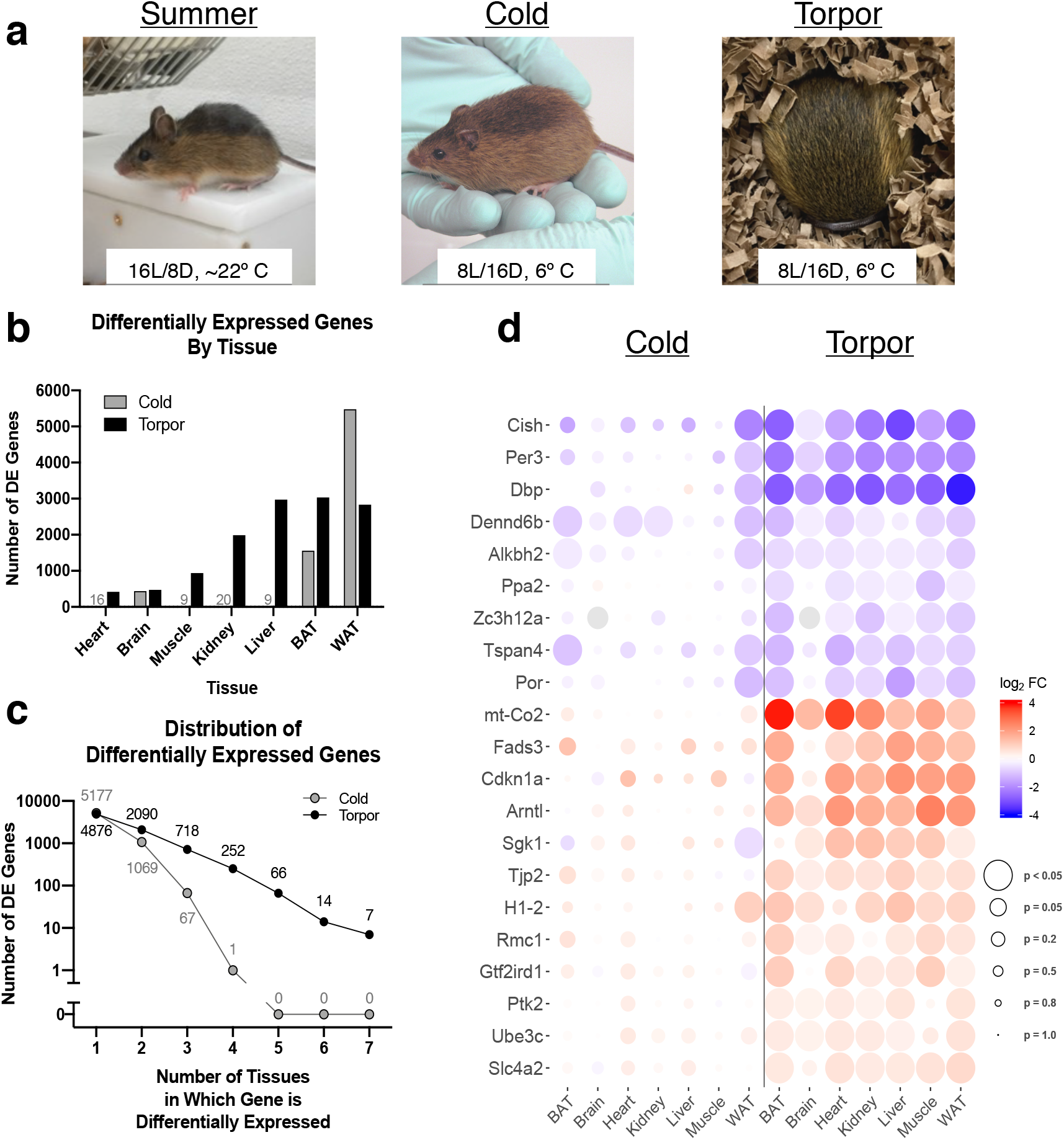
Gene expression in the meadow jumping mouse during fattening and hibernation. (**a**) Induction of fattening and hibernation by short photoperiod and cold temperature. Physical and housing conditions of “Summer,” “Cold” (fat, pre-torpor), and “Torpor” animals are shown. (**b**) Number of genes differentially expressed by tissue in meadow jumping mice that had prepared to hibernate in response to cold, short day conditions (“Cold”), and torpid animals (“Torpor”), relative to control animals under simulated summer conditions. (**c**) Few genes are differentially expressed in most tissues during torpor and preparation for hibernation. The number of genes is plotted according to the number of tissues in which each gene is differentially expressed. (**d**) Heatmap showing the change in expression, relative to control, of the 21 genes differentially expressed in 6 or 7 tissues during torpor. Color indicates limma^87^ log2-fold change; red and blue represent increased and decreased expression, respectively. Dot size is inversely correlated to adjusted *P*-value and provides a visual indication of statistical significance.

We performed differential expression analyses on the gene expression data from each tissue by comparing the ‘cold’ and ‘torpor’ conditions to the ‘summer’ control, with full results available in Tables S3-S9. In the ‘cold’ condition, BAT and WAT exhibited the largest number of differentially expressed genes (DEGs) relative to other tissues (Fig. 2b). During torpor, gene expression changes were more widespread, with kidney, liver, BAT, and WAT all having a relatively large number of DEGs (Fig. 2b). Each tissue type has significant overlap (*P*-values: 2.2×10^−108^, 3.3×10^−18^, 5.1×10^−8^, 1.1×10^−6^, 2.2×10^−6^, and 2.1×10^−10^ for BAT, brain, heart, kidney, muscle and WAT, respectively) between DEGs in the ‘cold’ and ‘torpor’ conditions, with the only exception of liver (*P*-value: 0.07), which has very few DEGs in the ‘cold’ condition. BAT and WAT had the greatest absolute number of DEGs shared between the ‘cold’ and ‘torpor’ conditions (Fig. S3).

### Universal DEGs across tissues for animals in torpor

The meadow jumping mouse experiences unique physiological demands during pre-hibernation fattening under cold exposure and during torpor. To better understand the holistic response of the organism to these conditions, we determined the distribution of DEGs across tissues (Fig. 2c). Consistent with the relatively low numbers of DEGs in tissues except BAT and WAT during pre-hibernation cold exposure, no genes were found to be differentially expressed in more than four tissues in the ‘cold’ condition. However, during torpor, a set of 7 genes had significant expression changes in all 7 tissues, and an additional 14 genes differentially expressed in 6 tissues (Fig. 2c). Among the 21 genes with differential expression in 6 or 7 tissues during torpor, the direction of the expression change (up- or downregulation) was analogous across all tissues for each gene (Fig. 2c, Table S10, suggesting a common regulatory response among tissues during torpor (Fig. 2d). The set of 21 genes encodes proteins with varied functions and some, including those with roles in circadian rhythm, membrane fluidity, and cell cycle arrest, are discussed below.

The endogenous circadian clock is driven by cycles of transcription and translation within a network of core clock genes including *Clock*, *Arntl*/*Bmal1*, and the cryptochrome and *Period* genes^24^. The core oscillator also depends on a transcriptional feedback loops involving Dbp, Rev-Erbα/β, and RORα/β as well as other inputs. Outputs from the endogenous circadian clock coordinate the molecular physiology and metabolism of individual tissues with daily rhythms of sleep, feeding, and activity. Evidence exists both for and against the maintenance of a circadian rhythm during hibernation in mammals^25^, and little is known about the molecular state of the endogenous clock during torpor. A study in European hamsters found that Per1, Per2, and Arntl/Bmal1 stopped oscillating in the suprachiasmatic nucleus during hibernation^26^.

We found that three of the 21 universal torpor DEGs were circadian genes. Averaging across all tissues, *Arntl*/*Bmal1* had a 29-fold increase in expression, and *Dbp* and *Per3* had 9.8-fold and 4.3-fold decreases in expression, respectively, during torpor relative to ‘summer’ controls (Fig. 2d). Despite being less significant statistically, expression of *Per1*, *Per2* and *Rev-Erbβ* (transcriptional inhibitor of *Arntl*/*Bmal1*) also decreased in all tissues by 1.3-fold, 2.6-fold, 2.2-fold on average, respectively. The expression level of other key players in circadian rhythm, including *Clock*, *Cry1*, *Cry2*, *Rev-Erbα*, and *RORα/β* fail to show a consistent pattern of expression changes across different tissues during torpor. There were no consistent differences in the expression of circadian clock genes in the ‘cold’ condition relative to ‘summer’ controls.

Notably, we found that the decreased expression of *Per3* is more prominent than *Per1* and *Per2* in torpid animals, implying that a reduced level of *Per3* might be more important for animals in hibernation. In contrast to Per1 and Per2, which collectively play an essential role in maintaining the daily circadian rhythm in mammals, Per3 does not seem to be essential in central circadian clock maintenance^27^. Loss of Per3 function affects adipogenesis and body mass accumulation in laboratory mice and genetic variations in Per3 are associated with type 2 diabetes in humans^28^. Taken together, the involvement of Per3 in regulation of sleep and metabolism with the observed significant decrease in *Per3* expression in hibernating jumping mice, suggest that Per3 may be an important factor in the biology of hibernation.

The large reduction in body temperature during torpor creates a challenge in maintaining fluidity in cell membranes. Homeoviscous adaptation, a strategy for increasing membrane fluidity in response to cold by desaturating membrane phospholipids, is employed by prokaryotes, lower eukaryotes, and some vertebrates including fishes^29^, but knowledge about its use by mammalian hibernators is limited^30,31^. Interestingly, the fatty acid desaturase Fads3, which exhibits widespread transcriptional upregulation during torpor (Fig. 2d), acts to desaturate sphingosine ceramides by adding a cis double bond to the sphingoid tail of the lipid^32,33^. Sphingolipids are important components of cell membranes and they have both structural and signaling roles in mammalian cells^34^. The upregulation of *Fads3* expression during torpor may serve to alter the signaling characteristics of sphingolipids or the physical properties of the membrane in response to torpor in the meadow jumping mouse. Fads3 shows elevated conservation in hibernating rodents compared to non-hibernating ones (discussed below), which further implies its important role in hibernation.

We observed upregulation of *Cdkn1a* in six of seven tissues during torpor (Fig. 2d). The protein product of *Cdkn1a*, p21, can act to inhibit cell cycle progression in proliferating cells and is induced in response to multiple stressful stimuli^35^. Increased expression of the cell cycle regulator p21 has been previously reported to induce cell cycle arrest in mildly hypothermic cells in culture^36^, and reports from mammalian hibernators have suggested the existence of a cell cycle arrest during torpor^37,38^. The widespread induction of *Cdkn1a* (p21) expression during torpor in the meadow jumping mouse suggests a role for this gene in response to the stressful conditions of torpor at the cellular level.

### Tissue-specific patterns of gene expression profile changes during torpor

To better understand the function of the large number of DEGs in individual tissues during fattening and torpor, we performed gene set enrichment analysis for each tissue using Gene Ontology (GO) annotations. The top three enriched GO terms in the categories of Biological Process, Cellular Component, and Molecular Function for each tissue are given by condition in Tables 1, S11, and S12, respectively. The most significant GO terms are plotted by tissue in Figures S4–S10 and complete results for all tissues are available in Table S13.

**Table 1.** Top three enriched Biological Process GO terms during fattening and torpor.

| **Tissue** | **Cold/Fattened** | **Torpor** |
| --- | --- | --- |
| **BAT** | *(no enrichment)* | GO:0009062 fatty acid catabolic process<br>GO:0072329 monocarboxylic acid catabolic process<br>GO:0032787 monocarboxylic acid metabolic process |
| **Brain** | *(no enrichment)* | GO:0016055 Wnt signaling pathway<br>GO:0198738 cell-cell signaling by wnt<br>GO:1905114 cell surface receptor signaling pathway involved in cell-cell signaling |
| **Heart** | *(no enrichment)* | GO:0045598 regulation of fat cell differentiation<br>GO:0071456 cellular response to hypoxia<br>GO:0072593 reactive oxygen species metabolic process |
| **Kidney** | GO:0016042 lipid catabolic process<br>GO:0044242 cellular lipid catabolic process<br>GO:2000036 regulation of stem cell population maintenance | *(no enrichment)* |
| **Liver** | *(no enrichment)* | *(no enrichment)* |
| **Muscle** | GO:0032787 monocarboxylic acid metabolic process<br>GO:0044283 small molecule biosynthetic process<br>GO:0072330 monocarboxylic acid biosynthetic process | GO:0055002 striated muscle cell development<br>GO:0055001 muscle cell development<br>GO:0051146 striated muscle cell differentiation |
| **WAT** | GO:0006325 chromatin organization<br>GO:0016071 mRNA metabolic process<br>GO:0016569 covalent chromatin modification | GO:0006631 fatty acid metabolic process<br>GO:0032787 monocarboxylic acid metabolic process<br>GO:0044283 small molecule biosynthetic process |

During torpor, DEGs in the brain were significantly enriched for annotations in the Wnt signaling pathway (Table 1, Fig. S5b). In the adult brain, Wnt signaling affects synaptic structure and plasticity and is important for maintenance of normal physiological function and adult hippocampal neurogenesis^39^. Inducible expression of Dkk1 (an antagonist of Wnt signaling) in transgenic mice results in reversible synapse loss in the adult hippocampus^40^. Hibernating ground squirrels exhibit significant, reversible reductions in the length and branching of dendrites and dendritic spines of hippocampal pyramidal cells^41^, and adult hippocampal neurogenesis is decreased in the hibernating hamster^42^, suggesting that Wnt signaling may play a role in these processes during torpor and arousal. Here, torpid meadow jumping mice exhibited differential expression in components of the canonical Wnt signaling pathway, including increased expression of *Dkk-1* (an antagonist), reduced expression of *Frizzled receptors 1 and 2*, and down-regulation of *Pygo1*, a positive regulator of b-catenin target gene expression (Fig. 3a and S11). These changes are broadly consistent with reduced Wnt signaling during torpor, suggesting a potential role for this signaling pathway in modulating synaptic activity and structure in the hibernating brain.

**Figure 3.**
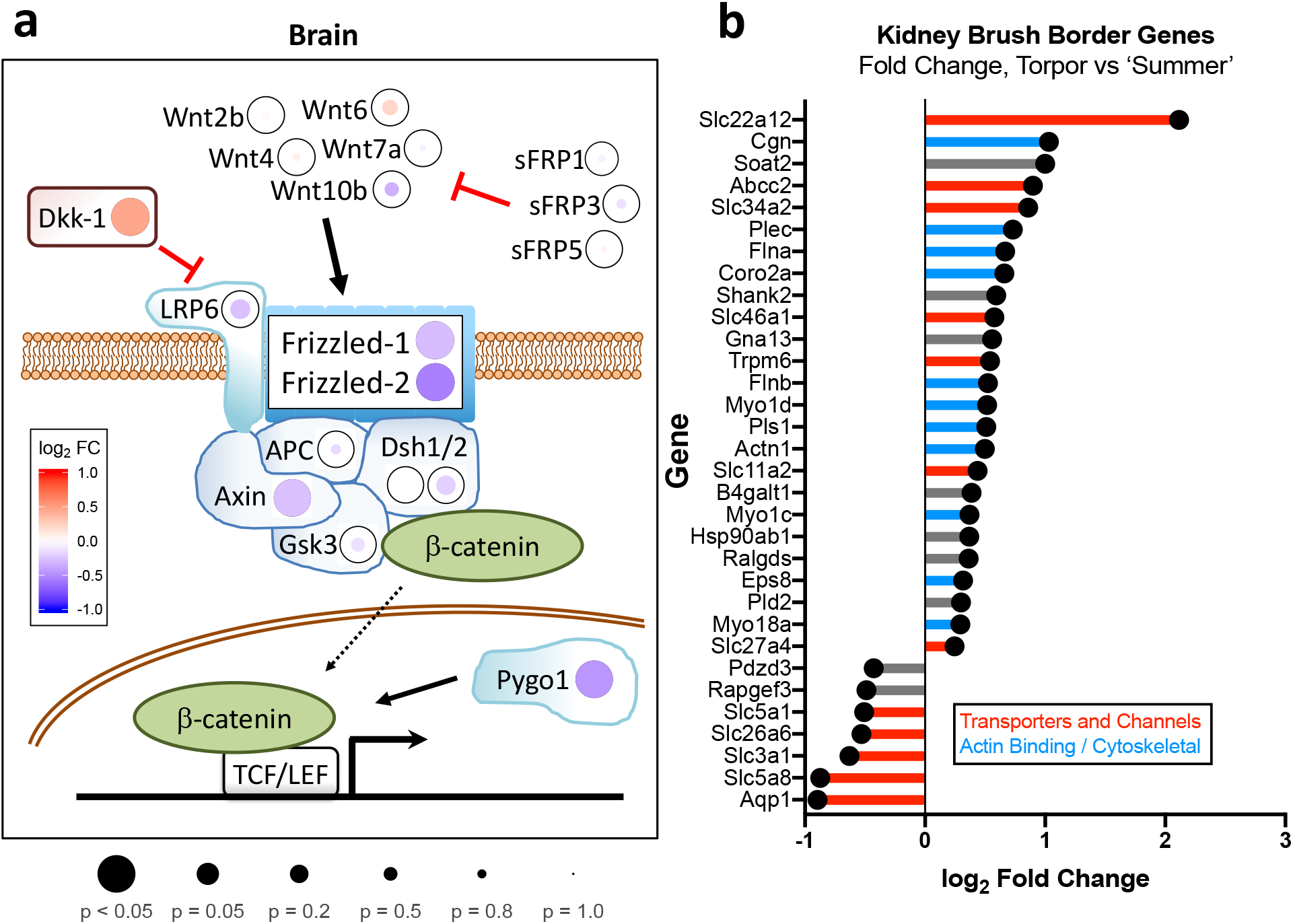
Gene ontology enrichment of differentially expressed genes reveals torpor -associated changes in brain and kidney. (**a**) During torpor, the brain exhibited enrichment in expression changes in the Wnt signaling pathway, including downregulation of Frizzled receptor and upregulation of the inhibitory factor Dkk-1. Color indicates log-fold change; red is up-regulation, blue is down-regulation. Dot size is inversely correlated to adjusted *P*-value and provides a visual indication of statistical significance. (**b**) The torpid kidney showed enrichment for expression changes in brush border genes. Genes with statistically significant (adj. p<0.05) expression changes and “brush border” GO annotation are plotted by log-fold expression change in torpor relative to “summer” control. The lines in the lollipop graph are colored according to manual annotation: red indicates membrane transporters and channels, and blue indicates proteins with actin-binding or other cytoskeletal function.

DEGs in the kidney during torpor tend to locate in the brush border (Table S11, Fig S7f). The brush border is found in the proximal tubules of the kidney, where microvillar structures on the luminal surface of the tubule increase surface area for reabsorption of water and solutes before they are lost in urine. Further investigation of these DEGs shows both up- and downregulation of water and solute transporters and upregulation of cytoskeletal proteins (Fig. 3b). The observed expression changes in genes encoding transporters is likely a consequence of the large change in kidney function during hibernation, as the animal ceases to eat and drink and must adjust renal function to maintain water balance and solutes normally filtered by the kidney^43^. Interestingly, the greatest change was found in urate transporter *URAT1* (*Slc22a12*) (Fig. 3b). Upregulation of this transporter during torpor may point to an increased need for urate reabsorption, possibly as a measure to maintain nitrogen balance during hibernation.

The effects of cold temperature on the kidney have been studied in the context of organ storage, during which the cold sensitivity of microtubules results in their destabilization and loss of cytoskeletal integrity at hibernation-like temperatures^44–46^. Cells from hibernating the thirteen-lined ground squirrels exhibit improved microtubule stability during cold exposure, and efforts to ameliorate cold-induced tissue damage in mouse kidneys during cold exposure link cytoskeletal stability to tissue health at low temperatures^47^. In the torpid meadow jumping mouse, the upregulation of genes encoding cytoskeletal proteins, including Cingulin, Plectin, Filamin-A/B, Plastin, and Alpha-actinin-1 (Fig 3b), may thus serve to support structural stability and function in the kidney at low temperature during hibernation.

### *Comparative analysis of* Z. hudsonius *and* Z. oregonus *genomes suggests potential mechanisms behind differential regulation of hibernation behavior*

We next sought to compare genomes of the meadow jumping mice (*Z. hudsonius*) with the Great Basin jumping mice (*Z. oregonus*). These two species are morphologically similar, yet they occupy different habitats and exhibit differences in how environmental cues such as photoperiod and food availability influence timing of breeding and hibernation. Because jumping mice speciation occurred relatively recently, we expected that the genetic variation within and between *Z. hudsonius* and *Z. oregonus* populations would provide clues as to the putative genetic bases of their behavioral differences and speciation. We therefore generated whole genome sequences of seven Great Basin jumping mice and six additional meadow jumping mice. The Great Basin jumping mouse samples were from the same subspecies previously used to study photoperiod control^20^ and were sampled from Utah and Wyoming in the United States of America^16^ (Fig. 4a, Table S14). The meadow jumping mouse samples were obtained from animals captured in Massachusetts (Fig. 4a). A summary of the relevant behavioral differences between the species and the relatedness of the individuals used in our comparative analysis is provided in Figure S2. We analyzed the protein sequences encoded by the genomes of both *Zapus* species and found that 915 genes were significantly divergent (F_ST_ > 0.8, and *P*-value < 0.05 for enrichment in divergent positions, Table S15), while 1079 genes showed evidence of positive selection (number of nonsynonymous substitutions / number of synonymous substitutions > 2 and *P*-value < 0.05, Table S16)^48,49^, and 414 genes were included in both categories (Fig. 4b, Table S17).

**Figure 4.**
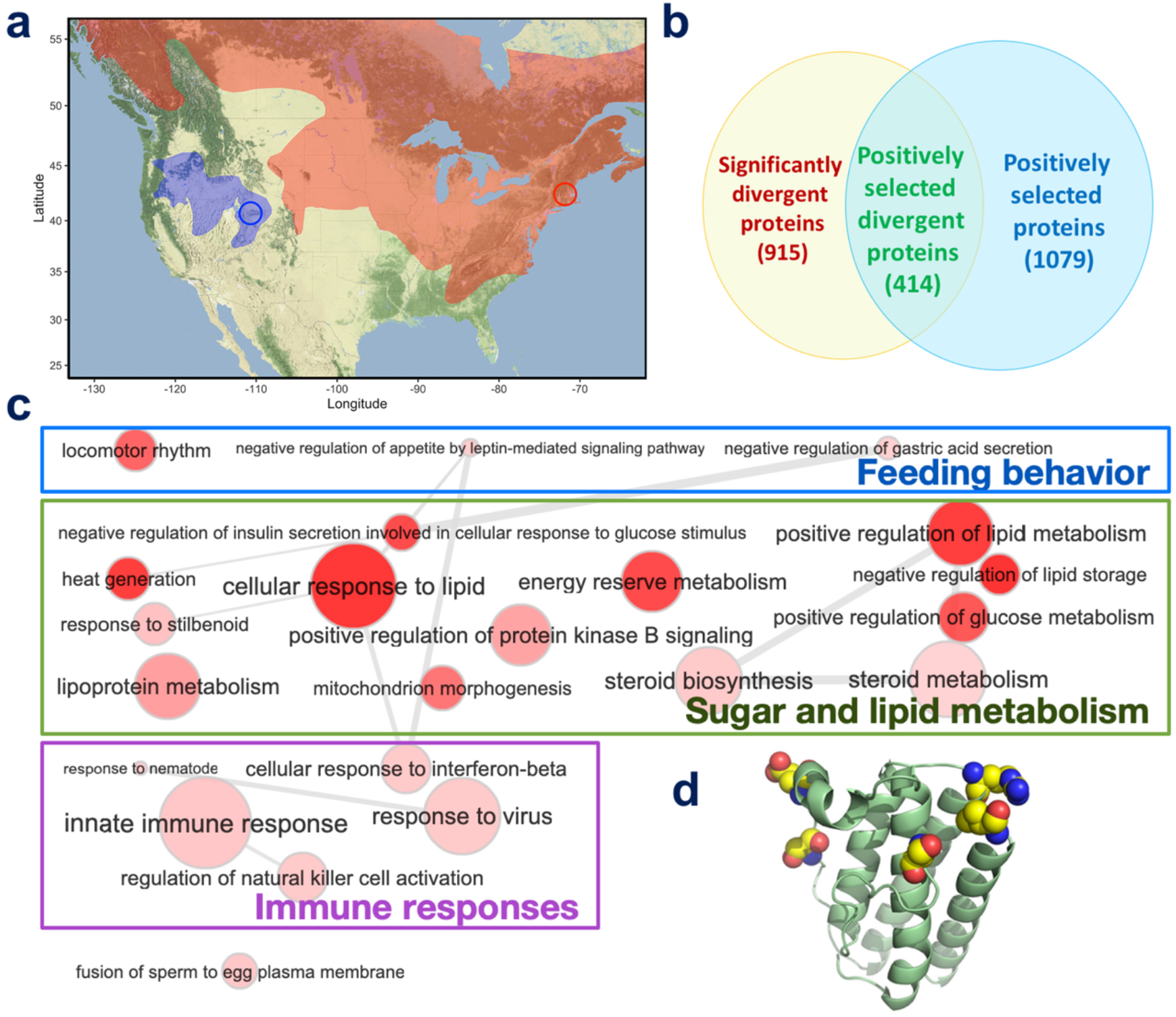
Comparative genomics of meadow (*Z. hudsonius*) and Great Basin jumping mice (*Z. oregonus*). (**a**) Geographic ranges of meadow (red) and Great Basin jumping mice (blue) with sampling locations for each marked with red and blue circles, respectively. (**b**) Venn diagram of genes significantly diverged and positively selected between the two species. (**c**) GO term enrichment for the set of positively selected and rapidly diverging proteins. Circle size indicates the number of genes associated with this GO term in *Mus musculus*, and intensity of red color visualizes statistical significance (darker color indicates lower *P*-value). Grey lines connect GO terms that tend to be associated with similar sets of proteins. (**d**) The significant divergence in leptin between the species is positively selected. The 3D structure model of leptin is shown as cartoon in green and atoms from amino acids that are different between the two species are shown as spheres colored by the atom type (yellow: carbon, green: nitrogen, red: oxygen).

The 414 genes that significantly diverged and were positively selected are potential drivers of speciation and thus are likely candidates responsible for phenotypic differences between the two *Zapus* species. We analyzed the function of these genes using GO terms (Table S18) and the significantly over-represented GO terms (false discovery rate < 0.1) can be classified into three areas: feeding behavior; sugar and lipid metabolism; and immune responses (Fig. 4c). Divergence in immune responses is reasonably expected because the two species were likely exposed to different pathogens as they evolved independently. However, divergence in feeding behavior and sugar/lipid metabolism may be an adaptation to different seasonal lifestyles. *Zapus hudsonius* does not fatten in presence of unlimited food supply when exposed to long photoperiods^8,18^, suggesting that they have a seasonal appetite-control mechanism linked to day length. In contrast, *Z. oregonus* starts fattening in presence of unlimited food supply regardless of day length^20^. This latter species has only a short active season in its native habitats^10^ and potentially does not need the same mechanism to control feeding behaviors. The difference in feeding and fattening behaviors may further indicate differential adaptive changes in sugar/lipid metabolism, which is required to properly channel the nutrients from food to either energy consumption or fat storage.

We found that the hormone leptin shows significant (*P*-value: 3.9×10^−6^) divergence between these two species and a strong signal of positive selection (*P*-value: 0.0027). Leptin has a key role in energy balance regulation and body weight control; however, the leptin receptor showed no evidence for rapid divergence or selective pressure, suggesting that the changes in leptin between *Zapus* species may have functional consequences. Leptin signaling controls appetite and decreases food intake and mutations resulting in loss of function cause over-feeding and obesity phenotypes in humans and mice^50,51^. Among the amino acid differences found in leptin (Fig. 4d), one polar to non-polar amino acid substitution (T48M) in *Z. oregonus* is of particular interest because mutation of the adjacent residue (H47N) results in an obesity phenotype in lab mice^52^. Consequently, the amino acid changes in leptin detected in *Z. oregonus* may decrease their ability to control appetite, and thus contribute to that species’ propensity to fatten when presented with an unrestricted diet^20^. In native habitats, the possibly reduced control of appetite might benefit *Z. oregonus* because they have a short active season and must rely on late-summer plant seed availability to store enough energy in fat tissue for prolonged hibernation of up to 9 months^53^. In contrast, *Z. hudsonius* controls appetite and fattening primarily in response to photoperiod, such that long summer days are permissive for an extended breeding season that lasts until fattening is triggered when day length shortens to near 12 hours in early autumn^18^.

### Comparison of rodent genomes

Using available rodent genomes, we performed multi-genome comparison with species classified as hibernating and non-hibernating (Table S19). Hibernation is an ancestral trait that was lost in animals that remain active throughout cold seasons^3^, and the ability to hibernate requires physiology and molecular machinery to function properly in two different conditions and to support the transition between physiological states. According to the genetics-physiology hypothesis, we predict that the set of proteins that play vital roles during torpor are likely to be more conserved in hibernating animals, while purifying selection on this same set of proteins may have decreased in species that lost the ability to hibernate. We therefore sought to identify proteins more conserved in hibernating rodents than non-hibernating ones (details in Methods).

This analysis yielded 171 proteins that met the significance threshold (*P*-value < 0.01, Table S20). We categorized the function of these proteins using the GO terms associated with them. The significantly (*P*-value < 0.01, Table S21) over-represented GO terms in the category of biological processes are summarized in Fig. 5a, and they belong to nuclear and cytosolic transport, cellular component assembly, cell adhesion, regulation of metabolism, and silencing of cell signaling. The enrichment of nuclear-cytosolic transport is the most significant and four proteins from the nuclear pore complexes (NPC) are among the proteins showing stronger conservation in hibernating animals. Regulation of nucleo-cytoplasmic trafficking is an important mechanism for plants to survive cold stress^54,55^, but the possible involvement of NPC or other nuclear and cytosolic transporting proteins are rarely studied in hibernating animals or in cold conditions. Considering that transcription is likely downregulated during hibernation, controlling the translocation of mRNA from nucleus to the cytosol may become an important mechanism to supply mRNA to sustain the expression of essential genes needed during hibernation.

**Figure 5.**
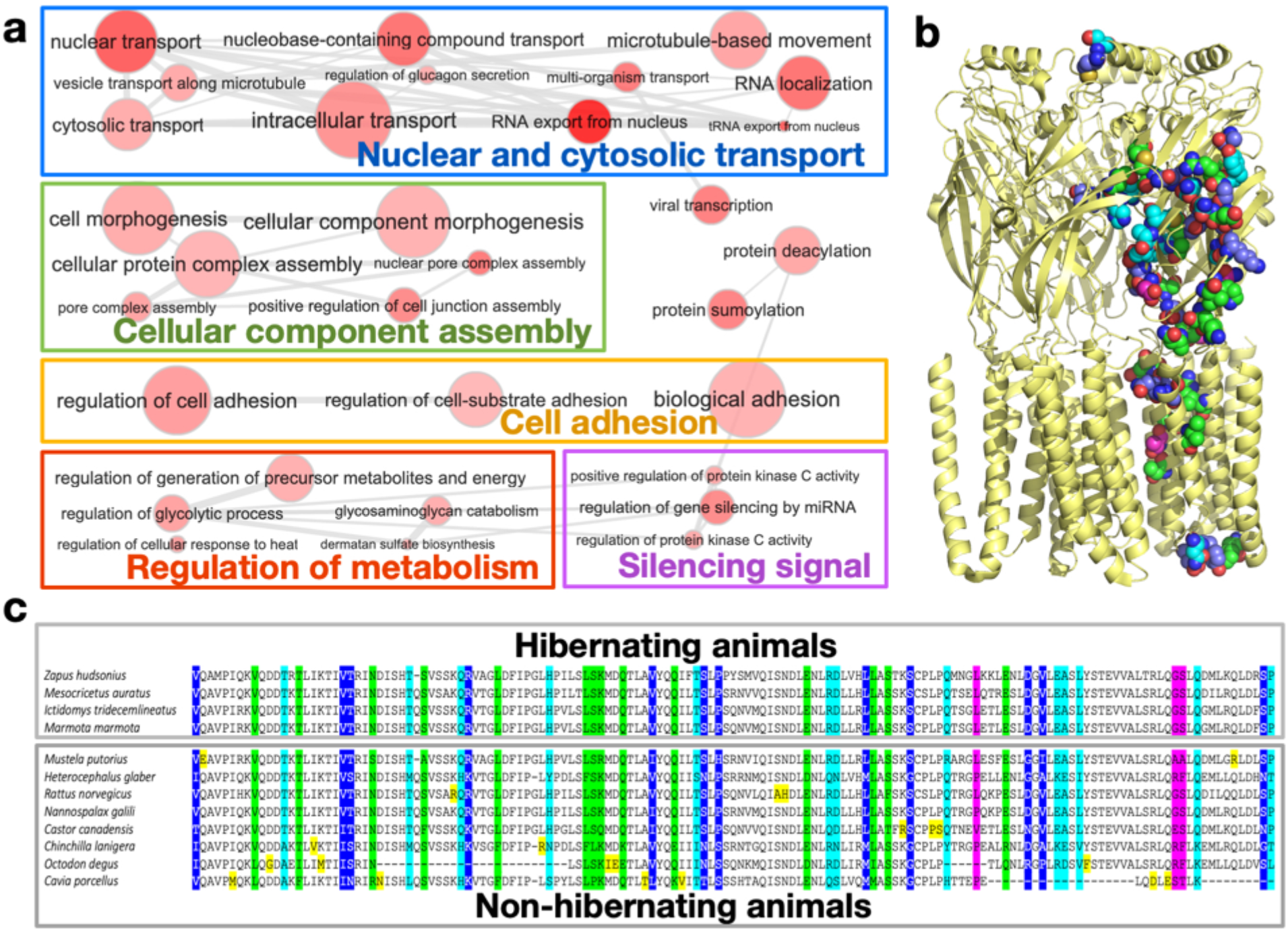
Proteins showing elevated conservation among hibernating rodents relative to non-hibernating mammals. (**a**) Gene ontology enrichment analysis of proteins enriched in positions that are conserved in hibernating rodents but variable in non-hibernating mammals. Circle size indicates the number of genes associated with this GO term in *Mus musculus*, and intensity of red color visualizes statistical significance (darker color indicates lower *P*-value). Grey lines connect GO terms that tend to be associated with similar sets of proteins. (**b**) The structure model of gamma-aminobutyric acid receptor (GABAR). GABAR subunit theta (GABRQ, modeled using the GABAR subunit beta2 as template) contains a high fraction of positions that are uniquely conserved in hibernating rodents, and atoms from residues in these positions are shown as spheres. Oxygen and nitrogen atoms are colored in red and blue, respectively. Carbon atoms are colored according to the conservation of this position in non-hibernating mammals: as the variability in non-hibernators increase, we color the carbons from green, through cyan and purple, to magenta. (**c**) Multiple sequence alignment of leptin from rodents representing different families (only *Ictidomys* and *Marmota* are from the same family). If a position only contains one mutation in non-hibernators, we color this mutation in yellow. If one position is mutated in 2, 3, 4, and >=5 rodents, we highlighted this position in green, cyan, blue, and magenta, respectively.

Proteins involved in cellular component assembly or inter-cell adhesion probably face stronger purifying selection in hibernating mammals. Although much of the normal enzymatic and cellular signaling activity is often reduced while hibernating, maintenance of proper cellular structure and interaction/adhesion between cells are likely in high demand during torpor. Temperature affects protein stability, affinity and cell adhesion^56,57^, and hibernating animals need to maintain proper affinity between molecular components of cells and extracellular matrices to preserve the structure and function of cells and tissues under a much wider temperature range than non-hibernators. This imposes additional physiological challenges to cellular maintenance genes in hibernating animals compared to the non-hibernating species, or those maintaining homeostasis.

Cell signaling and sensing activity are expected to be downregulated during torpor. We observed elevated conservation in regulators of protein-kinase C (PKC), such as Frizzled-2 and EGFR. PKC is involved in desensitization of G-protein couple receptors (GPCRs)^58^, and thus may play a role in reducing the activity of GPCRs in sensing the environmental and internal signals during hibernation. We also detected elevated conservation of gamma-aminobutyric acid receptor subunit theta precursor (GABRQ) in hibernating rodents (Fig. 5b). GABRQ is a subunit of the GABA receptor, and the activation of GABA receptors reduces the excitability of neurons which inhibits synaptic activity^59^. As suggested by the observed downregulation of Wnt signaling genes in the torpid brain, decreased synaptic activity during hibernation, coupled with activated GABA receptors, especially the theta subunit, may functionally inhibit brain activity during hibernation.

Finally, the uniquely conserved proteins in hibernating rodents include metabolic regulators that control glycolytic processes and energy generation. Hibernation requires a series of metabolic changes during the pre-hibernation fattening stage and during both torpor and arousal. Regulators that facilitate the switch between and maintenance of these extreme metabolic states are expected to be essential for hibernating animals. Among single-copy orthologs present in rodents, the metabolic hormone leptin has among the highest percentage of amino acid positions that are conserved in hibernators yet variable in non-hibernating congeners (Fig. 5c), indicating its importance in hibernation across mammals. For example, an investigation of leptins from hibernating and non-hibernating bats found both structural and functional differences in those proteins^60^. Leptin controls feeding behavior and energy balance, so it may also serve as an important feedback signal in at least some hibernating species because its levels increase as lipid stores accumulate in WAT before hibernation^61^. A more complete understanding of the role of leptin in pre-hibernation fat accumulation and subsequent switch to fat utilization requires additional lab and field-based experimental studies.

## MATERIALS AND METHODS

### Preparation of genomic DNA libraries and sequencing

For the reference assembly, DNA was obtained from the tail tissue of a wild-caught female meadow jumping mouse (*Z. hudsonius*) by proteinase-K digestion and phenol chloroform extraction. DNA from additional meadow jumping mice was obtained from ear tissue by spin-column purification. Prepared DNA samples from Great Basin jumping mice (*Z. oregonus*) were obtained from the Division of Genomics Resources at the Museum of Southwest Biology derived from wild-caught individuals^16^ and spin-column purification of heart or liver (Table S14).

For the reference assembly, the Broad Institute of MIT and Harvard prepared paired-end libraries and obtained 2×250 bp paired-end reads on the Illumina Hiseq2500 platform as previously described^62^. We prepared mate-pair libraries for the reference assembly with inserts of 3, 6, and 15kb as detailed elsewhere^63^. We prepared paired-end libraries for other samples using NEB Ultra II DNA library prep kit following our published protocols^64,65^. Paired-end libraries for additional genomes and mate-pair libraries for the reference assembly were sequenced on Illumina NextSeq and HiSeq 2500 instruments, respectively, in the UT Southwestern McDermott Center Next Generation Sequencing Core Facility. Hi-C libraries were generated and sequenced as a commercial service by Novogene using fresh frozen liver tissue from the animal used for reference genome assembly.

### Genome assembly and annotation

We assembled a reference genome from a *Zapus hudsonius* specimen as previously described^63,66^. All reads were processed by Trimmomatic^67^ to remove adapter sequences and low-quality bases (quality score < 20), and genomic DNA reads were further processed by Quake^68^ to correct sequencing errors. We used Platanus^21^ to assemble the genomes using the paired-end and mate-pair reads. Final assembly was performed using the 3D-DNA assembler and Juicebox Assembly Tools^23^ using the Hi-C reads provided by Novogene as a commercial service.

We used RepeatModeler <http://www.repeatmasker.org/RepeatModeler/> to identify repeats in the genome. In addition, since repeats with highly similar sequences may be erroneously combined into one in the genome assembly, we identified them using very high sequence depth (more than 4 times of the expected value) after mapping all the sequence reads to the draft genome using BWA^69^. We combined the repeats identified by RepeatModeler and our sequence depth criteria with repeats in Repbase^70^ to generate species-specific repeat libraries, and these libraries were supplied to RepeatMasker <http://www.repeatmasker.org/> to annotate repeats in the genome.

We annotated protein-coding genes in the genome using three approaches: homology-based, transcript-based, and *de novo* gene prediction. We used protein sets from *Cavia porcellus*, *Cricetulus griseus*, *Heterocephalus glaber*, *Ictidomys tridecemlineatus*, *Jaculus jaculus*, *Marmota marmota*, *Mesocricetus auratus*, *Mus musculus*, *Rattus norvegicus*, *Peromyscus maniculatus*, *Oryctolagus cuniculus* and *Homo sapiens* as references for homology-based annotation. The reference protein sets were aligned to the genome assembly using exonerate^71^. We aligned the RNA-seq reads from five tissues (brain, heart, liver, lung, and kidney) to the reference genome using TopHat ^72^, and derived transcript-based annotations using Cufflinks^73^. Three *de novo* gene prediction methods: Augustus^74^, GeneMark_ES^75^, and SNAP^76^ were used to obtain *de novo* gene annotations. We trained these *de novo* gene predictors for the *Zapus hudsonius* genome using confident gene models derived from the consensus between transcript-based and homology-based annotations. Finally, annotations by different approaches were combined in EvidenceModeler^77^ to obtain their consensus as the final gene predictions. We predicted the functions of these proteins by finding the closest homologs in Swissprot^78^ using BLASTP^79^ (E-value < 0.00001) and transferred the GO terms^80^ and functional annotations.

We assembled the genomes of other animals (including *Z. hudsonius* and *Z. oregonus*) by mapping the reads to the reference genome and SNP calling. We removed the sequencing adapters and low-quality portions from sequencing reads using Trimmomatic^67^, and merged read1 and read2 from the same fragment if their sequences overlap using PEAR^81^. The resulting reads of each specimen were mapped to the reference genome using BWA^69^. We kept reads that mapped unambiguously in the correct orientation from the BWA’s result. We performed SNP calling for each specimen using GATK^82^.

### Animal experiments

Animal work described in this manuscript has been approved and conducted under the oversight of the UT Southwestern Institutional Animal Care and Use Committee.

Core body temperature measurements were obtained from abdominal telemeter-loggers (nano-RFT, Star-Oddi, Iceland). Telemeters were surgically implanted in the abdomen. Briefly, aseptic surgery was performed under isofluorane anesthesia following administration of buprenorphine SR analgesic. The telemeter was placed in the abdomen through an incision in the skin at the midline and through the body wall in the linea alba. The body wall was sutured and skin closed with wound clips. All animals were given at least 2-weeks recovery before induction of pre-hibernation fattening or other manipulations.

Environmental conditions for the animals were as follows: ‘summer’ controls (16 hours light / 8 hours dark, 20°C), ‘cold’ and ‘torpor’ animals (8 hours light / 16 hours dark, 6°C). Light cycle and temperature were controlled using Powers Scientific environmental chambers (model RIS33SD).

Tissue harvest: non-torpid animals were euthanized by cervical dislocation following carbon dioxide narcosis; euthanasia of all non-torpid animals was performed within the 2 hours preceding the end of the light cycle. Torpid animals were euthanized by cervical dislocation approximately 72 hours into their 2^nd^ or 3^rd^ torpor bout, with the expectation that this would be in the middle of the torpor bout as previously observed^8^. Onset of hibernation and torpor bouts were monitored via body temperature telemetry, supplemented in some cases by passive IR motion detectors and nest box temperature as previously described^8,83^. Following euthanasia, tissues were collected rapidly and in a standardized order for each animal. Samples for RNA analysis were preserved in *RNALater* and frozen at −80°C.

### Transcriptome sequencing and analysis of differentially expressed genes

RNA samples were extracted from *RNALater* preserved tissues using TriZol reagent according to manufacturer’s protocol. Carrier glycogen was added to white adipose tissue samples to facilitate recovery of RNA. RNA integrity was determined via TapeStation RIN scores, and high-quality RNA was used to generate directional mRNA libraries. Total RNA was DNase treated, mRNA was enriched using NEBNext Poly(A) mRNA isolation beads (NEB #E7490) and libraries were assembled using the NEBNext Ultra II Directional RNA Library Prep Kit for Illumina (NEB #E7760) with NEBNext Multiplex Oligos for Illumina primer sets (NEB #E7335 and #E7500). mRNA libraries were quality controlled via TapeStation then pooled and sequenced using single-end 75 bp reads on an Illumina MiSeq instrument in the UT Southwestern McDermott Center Next Generation Sequencing Core Facility.

RNA sequencing reads were analyzed using the HISAT, StringTie and Ballgown pipeline^84^, then mapped to the *Mus musculus* proteins in the Uniprot database using TBLASTN^85^. Further analysis was performed using R^86^. Gene expression for each tissue was visualized using histograms generated in R, and low expression genes were filtered for each tissue by excluding genes with summed expression across all conditions smaller than 10; this method of filtering retains genes that are expressed in some conditions but not others. The limma package^87^ was used to perform differential expression analysis on the filtered data using a one-factor analysis; that is, the ‘cold’ and ‘torpor’ conditions were compared to ‘summer’ controls for each tissue. Results tables from limma were generated using Benjamini & Hochberg (“BH”) multiple hypothesis correction and genes were determined to be significantly differentially expressed using a q-value (adjusted *P*-value) cutoff of 0.05. Genes that were significantly differentially expressed in the ‘cold’ and ‘torpor’ conditions for each tissue were visualized using the VennDiagram package^88^. The clusterProfiler package was used to perform GO analysis and visualization^89^. Heatmaps were produced in R using ggplot2^90^; circle size was plotted based on a transformation of adjusted *P*-value to provide a visual representation of statistical significance^91^, but with a step in size at the chosen significance level of 0.05. The *Zapus* range map was generated using rgdal^92^ and ggmap^93^ in R.

### *Comparative analysis of* Z. hudsonius *and* Z. oregonus

In order to identify proteins showing significant divergence between the two *Zapus* species, we first computed the fixation indices (F_ST_) for each protein. F_ST_ is calculated as 1 - π_within_/π_between_, where π_within_ and π_between_ are the average divergence between a pair of animals from the same and different species, respectively. We required divergent proteins to show F_ST_ higher than 0.8. A high F_ST_ indicates that the protein is relatively conserved within a species, thus helps to rule out proteins intrinsically variable due to the lack of important function. Second, we detected all the positions conserved (sharing a common amino acid in over 80% haplotypes) within but different between species, and further identified proteins significantly enriched (*P*-value < 0.05) in such positions. Enrichment was quantified using binomial tests (p = rate of divergent positions summarized over all proteins, m = the number of divergent positions in a protein, n = the total number of aligned positions in a protein).

We tested whether a gene was positively selected during the history of divergence between species using the rate of nonsynonymous mutations. For each gene, we estimated the number of nonsynonymous substitutions (N_NS_) and synonymous substitutions (N_SS_) needed to change from the dominant (present in >60% haplotypes) codon of one species to the dominant (present in >60% haplotypes) codon of another species. If there were multiple substitution paths to change from the codon of *Z. hudsonius* to the codon of *Z. oregonus*, the path with the fewest number of nonsynonymous substitutions (parsimonious) was taken. We summarized the numbers over all genes to obtain the total numbers of synonymous (TN_SS_) and nonsynonymous substitutions (TN_NS_) and computed the average nonsynonymous mutation rate as R_aver_ = TN_SS_/(TN_SS_+TN_NS_). To calculate the statistical significance for positive selection in each gene, we used binomial tests to compare if N_NS_ / (N_NS_ + N_SS_) was significantly larger than R_aver_. A gene with *P*-value less than 0.05 was considered to be positively selected.

For genes that show both a sign of positive selection and encode significantly diverged proteins between the two species, we analyzed their function using GO terms. Enriched GO terms associated with this set of genes were detected using binomial tests (m = the number of proteins in this set that were associated with this GO term, N = number of proteins in this set, p = the probability for this GO term to be associated with any protein encoded by the *Zapus* genome). We further corrected the statistical significance for multiple tests and computed the false discovery rate (q-value) for each enriched GO term using “BH” procedure^94^. GO terms with q-values lower than 0.05 were visualized using REVIGO^95^. Interesting examples among these positively selected divergent proteins were studied manually. We searched for homologous 3D structures of these proteins in the Protein Data Bank using BLASTP, and the BLAST alignment was used to find the corresponding residue in the 3D structure for each residue in the query protein.

### Comparison of hibernating and non-hibernating rodents

We obtained the alignments of orthologous proteins in mammals from OrthoMaM database. We selected one representative species per rodent family from the available organisms in this database, and collectively sampled each of the four major clades of rodents (Hystricomorpha, Sciuromorpha, Castorimorpha, Myomorpha) plus a carnivore. We excluded rodents that can undergo torpor but do not normally hibernate over the entire winter, and the resulting species for the analysis and the families they represent (in parenthesis) are as follows: *Mesocricetus auratus* (Cricetidae, hibernator), *Ictidomys tridecemlineatus* (Sciuridae, hibernator), *Marmota marmota* (Sciuridae, hibernator), *Nannospalax galili* (Spalacidae), *Heterocephalus glaber* (Heterocephalidae), *Rattus norvegicus* (Muridae), *Octodon degus* (Octodontidae), *Castor canadensis* (Castoridae), *Cavia porcellus* (Caviidae), *Chinchilla lanigera* (Chinchillidae), and *Mustela putorius* (Mustelidae). The first three species (two families) undergo hibernation and are the only representative hibernating rodents available in OrthoMaM. We therefore used both *Ictidomys tridecemlineatus* and *Marmota marmota* from Sciuridae.

From the multiple sequence alignments of the above species, we removed positions dominated by gaps. We derived the consensus sequence for each orthologous group by taking the most frequent amino acid in each position. The consensus sequence was used to search against the transcript sequences we obtained for *Z. hudsonius* with TBLASTN, and the best hit was added to the multiple sequence alignment. We thus obtained multiple sequence alignments for 13,031 single-copy orthologous genes from 3 hibernating rodent families and 8 non-hibernating mammals (7 rodent families).

We took positions containing non-gap residues in all three hibernating rodent families and at least three non-hibernators for the following analysis. We identified the positions where the three hibernating rodent families contain identical amino acids and counted the fraction of non-hibernators showing different amino acids at this position. This fraction was summed over all positions and rounded to an integer to reflect the average number of mutations (N_mut_) in non-hibernating mammals at conserved position for hibernating rodents. Proteins containing a high fraction (normalized by the total number of positions, N_total_) of such mutations may either be more conserved among hibernating rodents for functional relevance or simply represent the more variable proteins where three selected families, even if randomly selected, may accidently possess the same amino acids due to mutation saturation. To rule out the latter scenario, we generated randomized controls by randomly selecting three families and counting the average number of mutations in other families at positions where the three random families contain identical amino acids. This average number of mutations was normalized by the total number of positions that are non-gap for the three selected families and at least 3 out of the 8 remaining families to obtain the mutation rate in a random control. We enumerated all possible random sets of three selected families (cannot be the three hibernating families) and computed the average rate, R_control_.

If the observed mutation rate (N_mut_ / N_total_) for comparing hibernating rodents against non-hibernators in a protein is significantly higher than R_control_, we considered the protein to show elevated conservation among the hibernating species. We evaluated the statistical significance using binomial tests and detected 171 proteins showing *p* value less than 0.1. The functional enrichment of these proteins and interesting cases among them were studied as described in the previous section.

## ACKNOWLEDGEMENTS

The authors thank Alyssa D. McNulty, Nick Grishin, James Cook, the Division of Genomic Resources at the Museum of Southwestern Biology, the UT Southwestern McDermott Center Next Generation Sequencing Core Facility, and the Texas Advanced Computing Center at the University of Texas at Austin for assistance and support. We thank Diane Genereux for helpful comments on the manuscript. This work was supported by the Sara and Frank McKnight Fund for Biochemical Research and NIH grant DP5OD021365 to WJI.

## AUTHOR CONTRIBUTIONS

W.J.I. and Q.C. conceived and co-lead the study. J.L.M. collected and prepared *Zapus oregonus* samples. E.A.B. and W.J.I. performed hibernation experiments. E.A.B. and W.J.I. prepared RNA sequencing libraries. Q.C. prepared genome sequencing libraries and W.J.I. assisted. J.A., J.J, E.K.K and K.L-T. sequenced the reference genome. Q.C. performed data preparation, genome assembly, transcriptome assembly, annotation, and comparative analyses. J.Z. performed computational analysis. W.J.I. performed RNA sequencing analysis and Hi-C genome assembly. W.J.I. and Q.C. wrote the paper with input from all authors.

## SUPPLEMENTAL FIGURES

**Supplemental Figure 1.**
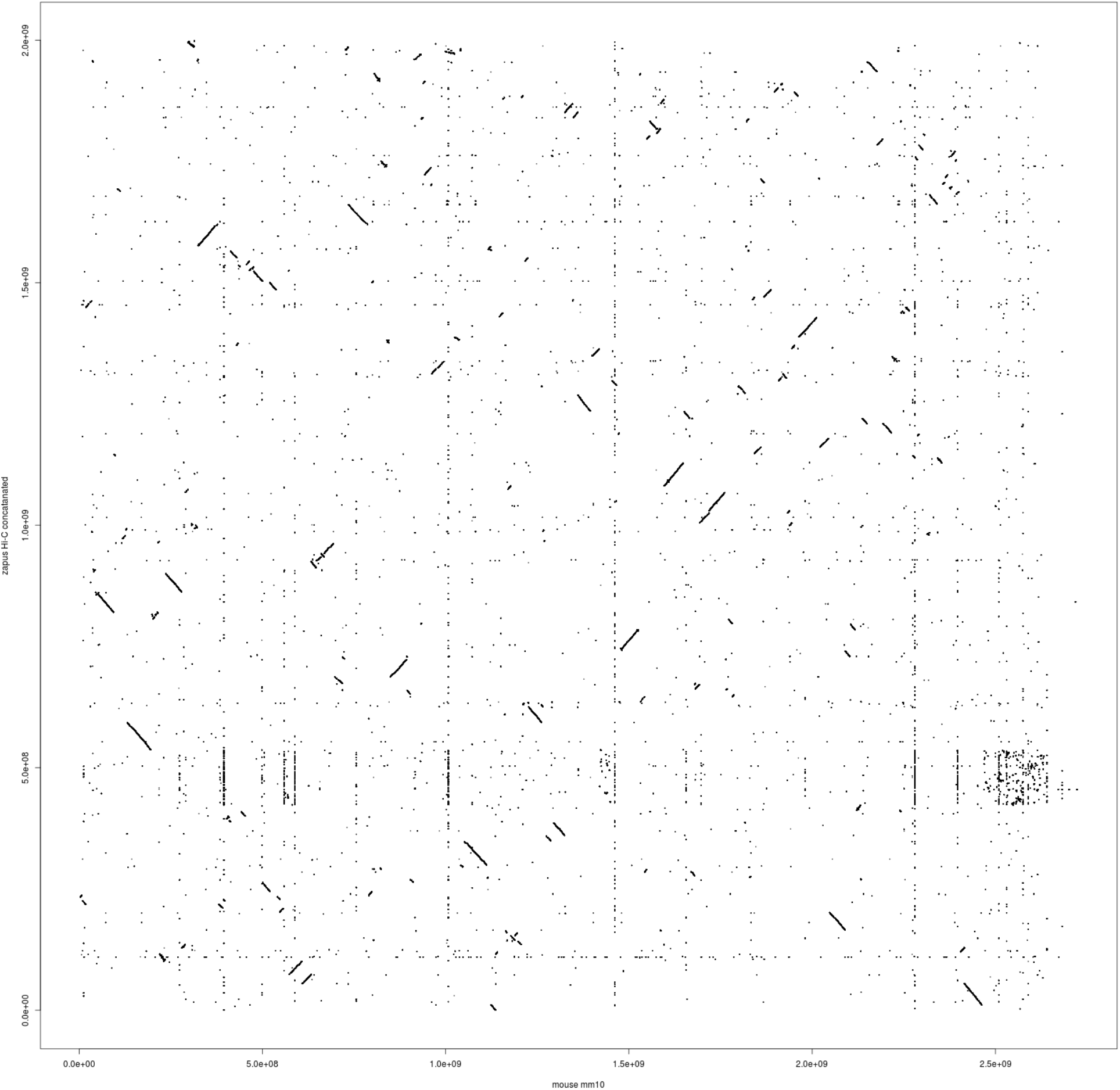
LASTZ alignment of concatenated *Mus* (mm10) and *Z. hudsonius* genome assemblies shows areas of synteny.

**Supplemental Figure 2.**
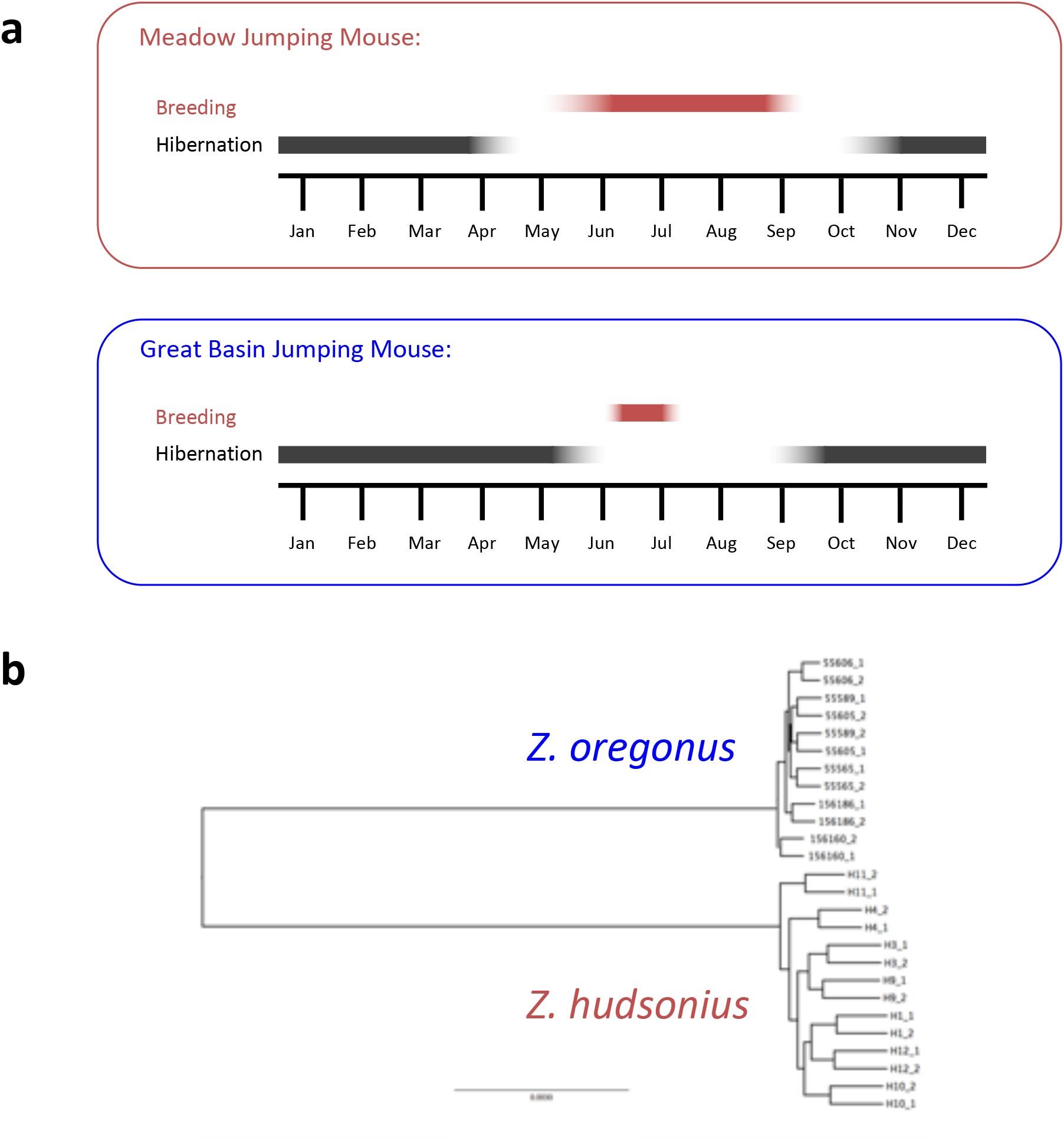
Stereotyped seasonal activity and relatedness of *Z. hudsonius* and *Z. oregonus* used in this study. (**a**) Meadow jumping mice (*Z. hudsonius*) exhibit a summer breeding season that allows multiple litters, while Great Basin jumping mice (*Z. oregonus*) breed once upon emergence from hibernation and have a shorter active season at higher altitudes from whence the samples originated. (**b**) Phylogenetic relationships of the *Zapus* genomes used in this study.

**Supplemental Figure 3.**
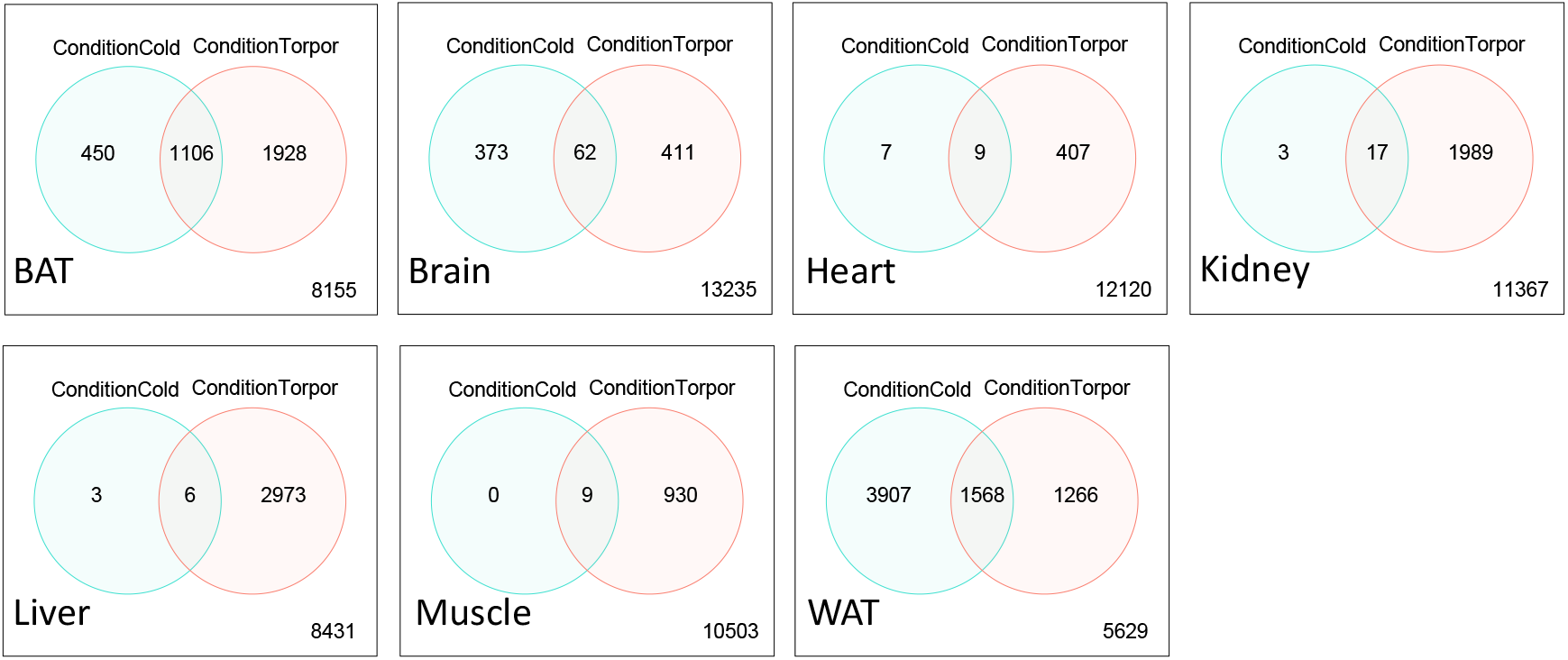
Overlap, by tissue, of genes differentially expressed in the ‘Cold’ and ‘Torpor’ condition. Venn diagrams showing the overlap of each set of differentially expressed genes in ‘Cold’ animals (pre-hibernation animals that had fattened in response to housing at 8L/16D photoperiod and 6°C) and ‘Torpor’ animals that were harvested after approximately ~72 hours of torpor, relative to lean animals held under simulated summer conditions. White and brown adipose show the greatest absolute number of differentially expressed genes with a large number of common changes.

**Supplemental Figure 4.**
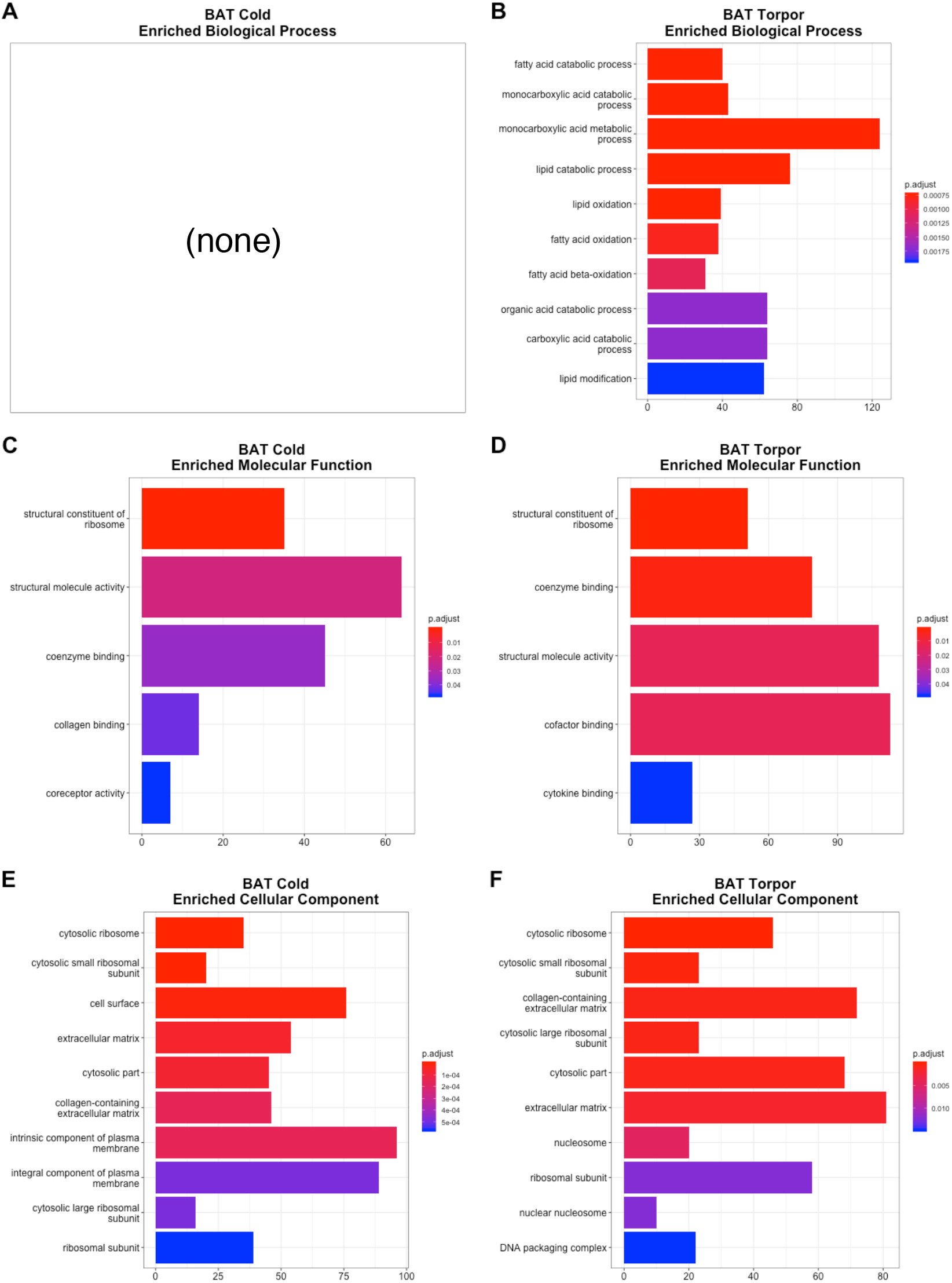
Brown Adipose Tissue: significantly enriched sets of differentially expressed genes by Gene Ontology term. Enriched ‘Biological Process’ terms for (**a**) ‘Cold’ and (**b**) ‘Torpor’ conditions. Enriched ‘Molecular Function’ terms for (**c**) ‘Cold’ and (**d**) ‘Torpor’ conditions. Enriched ‘Cellular Compartment’ terms for (**e**) ‘Cold’ and (**f**) ‘Torpor’ conditions.

**Supplemental Figure 5.**
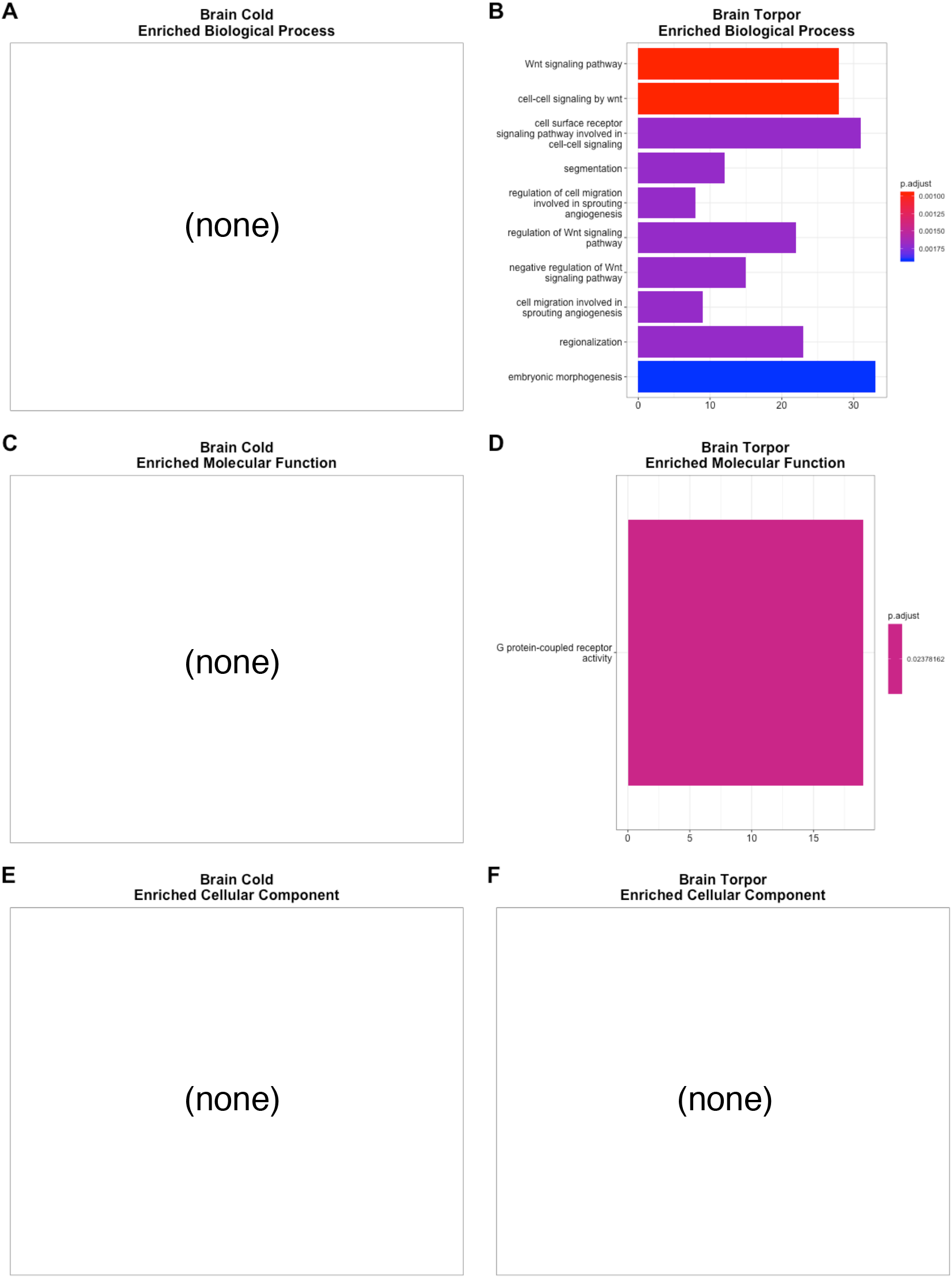
Brain: significantly enriched sets of differentially expressed genes by Gene Ontology term. Enriched ‘Biological Process’ terms for (**a**) ‘Cold’ and (**b**) ‘Torpor’ conditions. Enriched ‘Molecular Function’ terms for (**c**) ‘Cold’ and (**d**) ‘Torpor’ conditions. Enriched ‘Cellular Compartment’ terms for (**e**) ‘Cold’ and (**f**) ‘Torpor’ conditions.

**Supplemental Figure 6.**
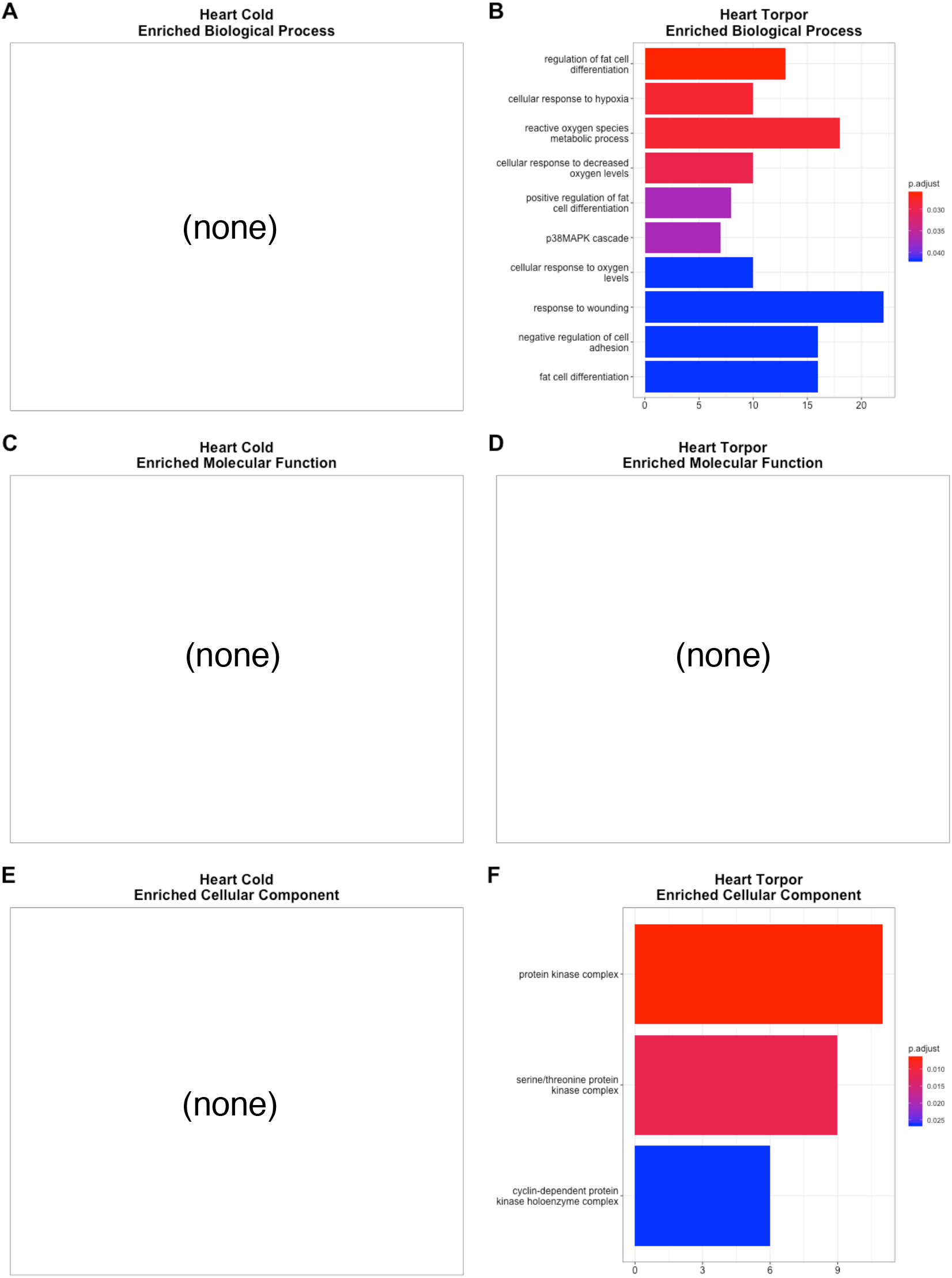
Heart: significantly enriched sets of differentially expressed genes by Gene Ontology term. Enriched ‘Biological Process’ terms for (**a**) ‘Cold’ and (**b**) ‘Torpor’ conditions. Enriched ‘Molecular Function’ terms for (**c**) ‘Cold’ and (**d**) ‘Torpor’ conditions. Enriched ‘Cellular Compartment’ terms for (**e**) ‘Cold’ and (**f**) ‘Torpor’ conditions.

**Supplemental Figure 7.**
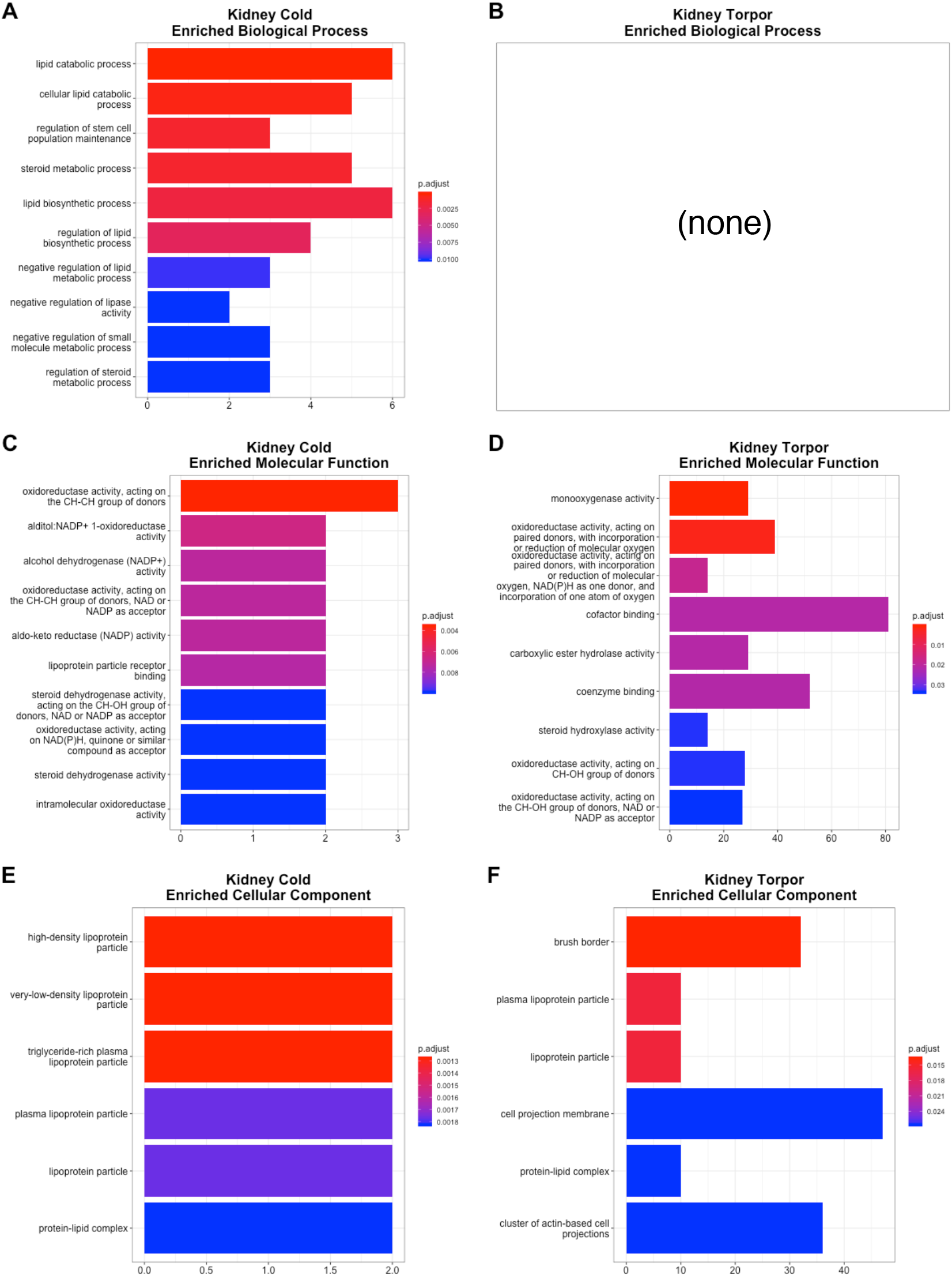
Kidney: significantly enriched sets of differentially expressed genes by Gene Ontology term. Enriched ‘Biological Process’ terms for (**a**) ‘Cold’ and (**b**) ‘Torpor’ conditions. Enriched ‘Molecular Function’ terms for (**c**) ‘Cold’ and (**d**) ‘Torpor’ conditions. Enriched ‘Cellular Compartment’ terms for (**e**) ‘Cold’ and (**f**) ‘Torpor’ conditions.

**Supplemental Figure 8.**
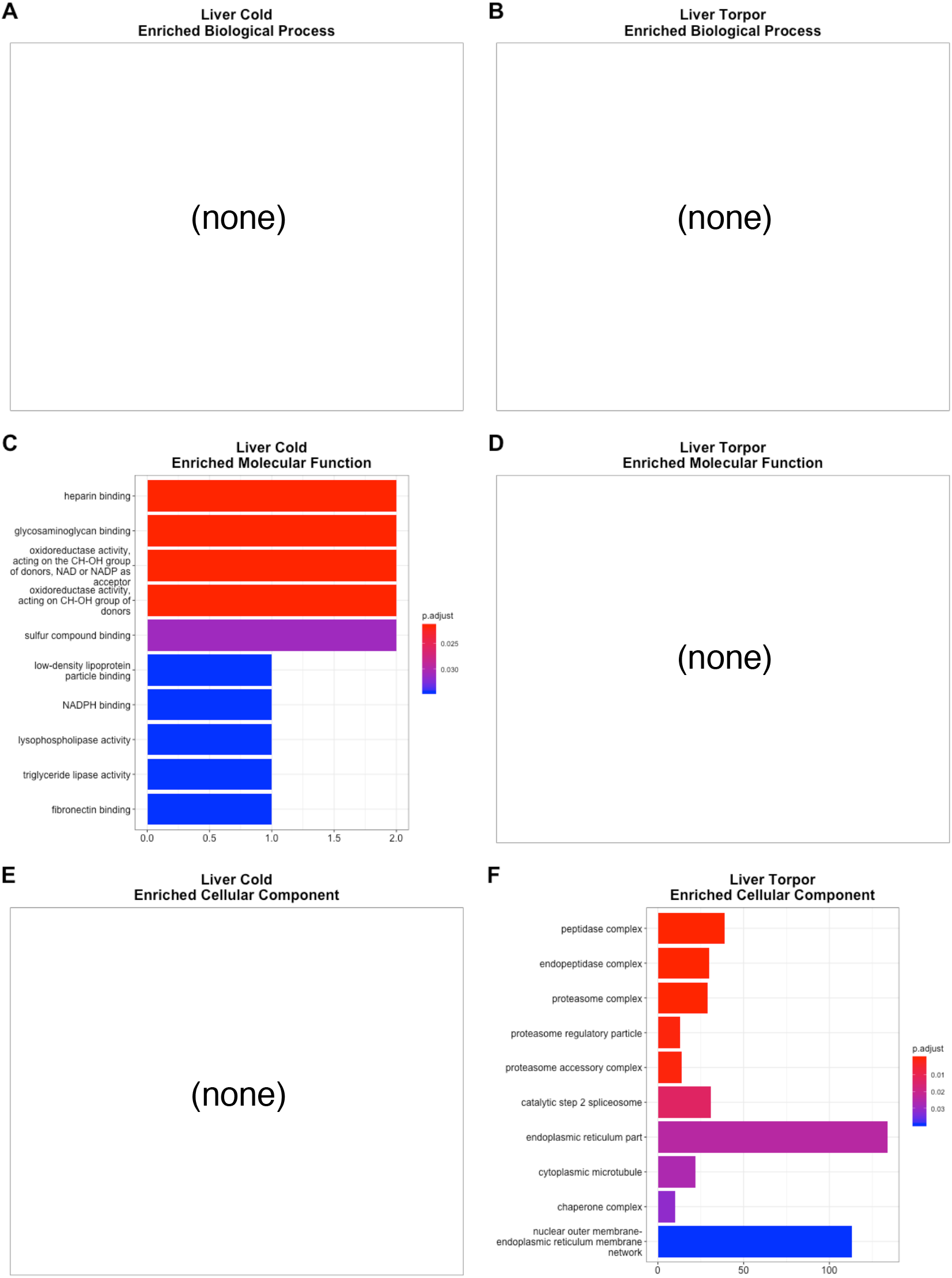
Liver: significantly enriched sets of differentially expressed genes by Gene Ontology term. Enriched ‘Biological Process’ terms for (**a**) ‘Cold’ and (**b**) ‘Torpor’ conditions. Enriched ‘Molecular Function’ terms for (**c**) ‘Cold’ and (**d**) ‘Torpor’ conditions. Enriched ‘Cellular Compartment’ terms for (**e**) ‘Cold’ and (**f**) ‘Torpor’ conditions.

**Supplemental Figure 9.**
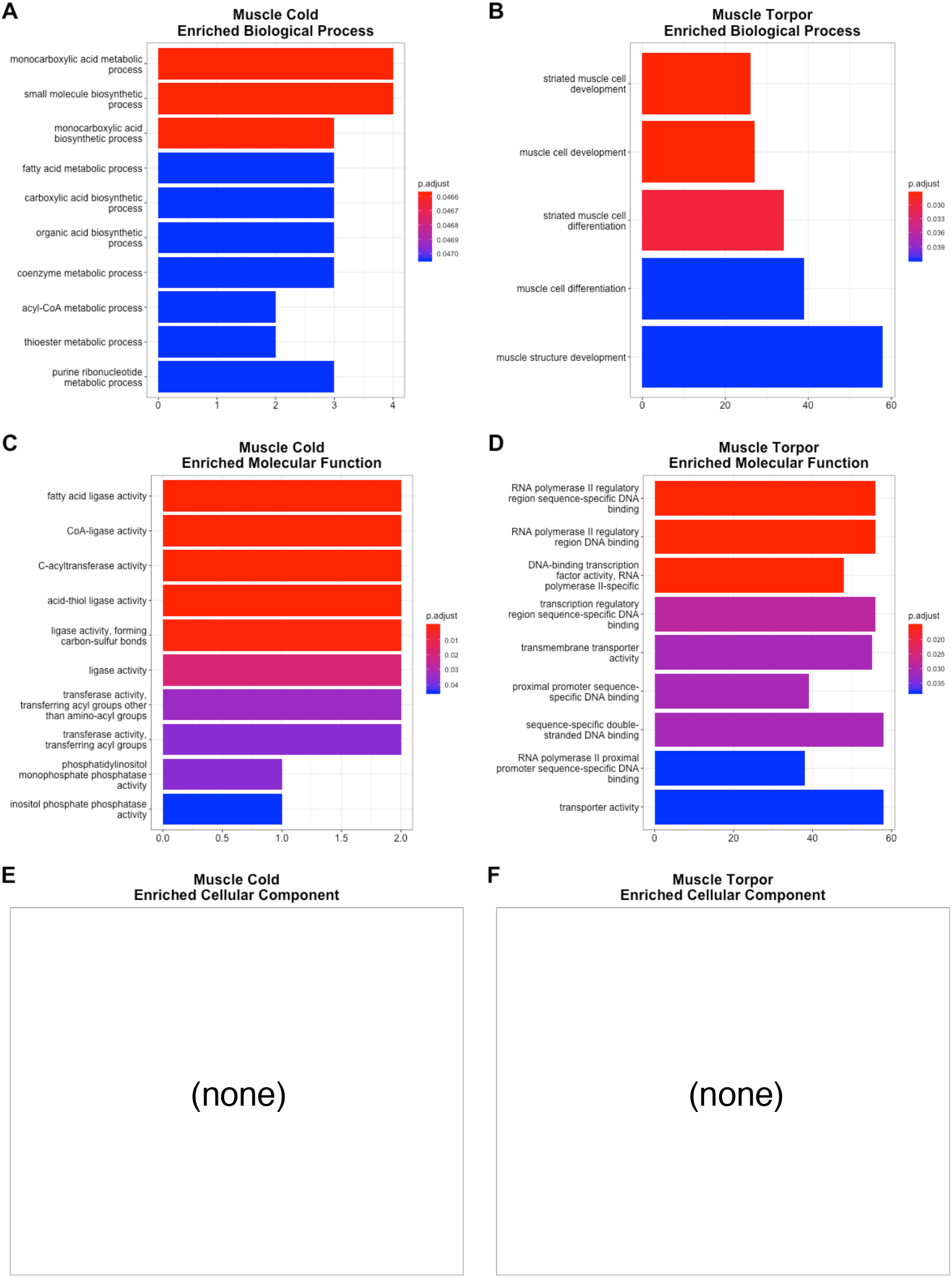
Muscle: significantly enriched sets of differentially expressed genes by Gene Ontology term. Enriched ‘Biological Process’ terms for (**a**) ‘Cold’ and (**b**) ‘Torpor’ conditions. Enriched ‘Molecular Function’ terms for (**c**) ‘Cold’ and (**d**) ‘Torpor’ conditions. Enriched ‘Cellular Compartment’ terms for (**e**) ‘Cold’ and (**f**) ‘Torpor’ conditions.

**Supplemental Figure 10.**
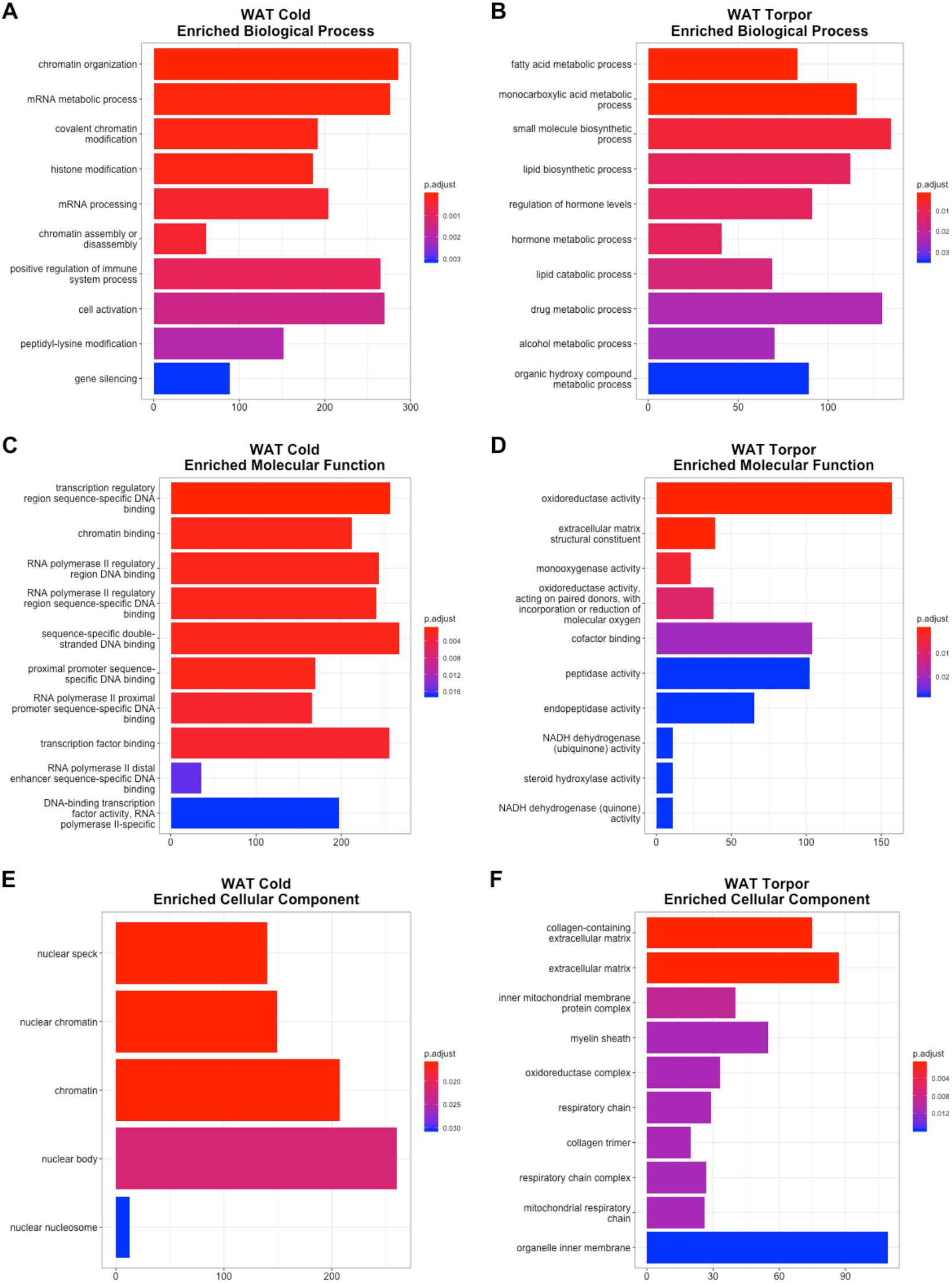
White Adipose Tissue: significantly enriched sets of differentially expressed genes by Gene Ontology term. Enriched ‘Biological Process’ terms for (**a**) ‘Cold’ and (**b**) ‘Torpor’ conditions. Enriched ‘Molecular Function’ terms for (**c**) ‘Cold’ and (**d**) ‘Torpor’ conditions. Enriched ‘Cellular Compartment’ terms for (**e**) ‘Cold’ and (**f**) ‘Torpor’ conditions.

**Supplemental Figure 11.**
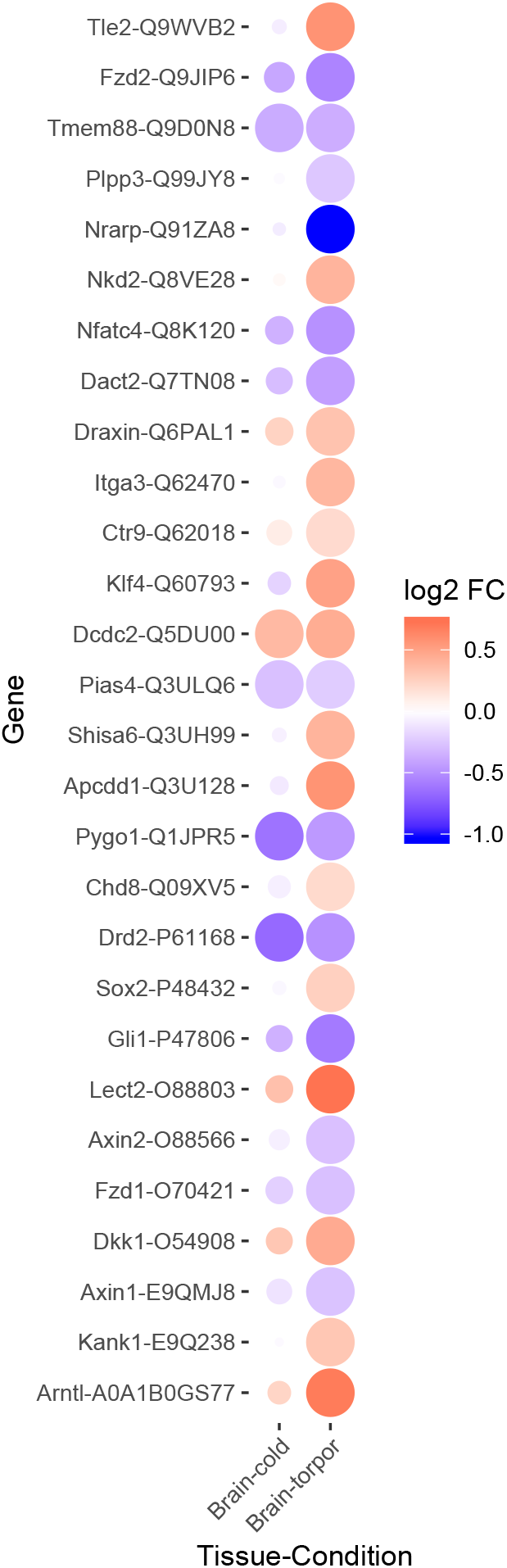
Heatmap of Wnt pathway gene expression changes in brain from ‘Cold’ and ‘Torpor’ animals. Color indicates log fold change; red is up-regulation, blue is down-regulation. The size of the dot is inversely correlated to adjusted *P*-value and provides a visual indication of statistical significance. The common gene name and corresponding mouse (*Mus musculus*) UNIPROT ID are indicated.

